# Submitochondrial Protein Translocation in Thermogenic Regulation

**DOI:** 10.1101/2023.05.04.539294

**Authors:** Fahrettin Haczeyni, James M. Jordan, Benjamin D. Stein, Sandra Steensels, Le Li, Vincent Dartigue, Selenay S. Sarklioglu, Jixuan Qiao, Xi K. Zhou, Andrew J. Dannenberg, Neil M. Iyengar, Haiyuan Yu, Lewis C. Cantley, Baran A. Ersoy

## Abstract

Mitochondria-rich brown adipocytes dissipate cellular energy as heat. Excessive nutrition or prolonged cold exposure suppress thermogenesis, but the mechanisms remain incompletely understood. Here we report that excessive cold or metabolic stress-induced proton leak into the matrix interface of mitochondrial innermembrane (IM) mobilizes 73 proteins from IM into matrix, which in turn alter mitochondrial bioenergetics. Interactome analysis indicates that key subunits of the electron transport chain and the mitochondrial calcium uniporter provide histidine-rich IM docking sites for 40 translocating proteins via pH-dependent protein-protein interactions. We further determine that 34% of translocating proteins correlate with obesity in human subcutaneous adipose tissue among which the top factor, acyl-CoA thioesterase 9 (ACOT9), enzymatically deactivates and prevents the utilization of acetyl-CoA in thermogenesis and promotes obesity. Overall, this study introduces stress-induced submitochondrial protein translocation as a new mitochondrial mechanism.

**One-Sentence Summary:** Mitochondrial stress regulates energy utilization by forcing translocation of IM-bound proteins into the matrix.

## Introduction

Brown and beige adipose tissue (BAT) do not only maintain body temperature^1–3^ but also provide a metabolic sink for the catabolic consumption of excess lipids and glucose^4–7^. Short-term cold exposure or acute bout of excessive food intake promote thermogenesis^8,9^, however, long-term obesogenic diet or prolonged cold exposure have complex effects on BAT physiology, which impedes thermogenesis^10–12^. While active downregulation of heat production, such as torpor or hypothermia, is an evolutionarily conserved mechanism for the conservation of limited energy sources^13–16^, it is unclear how chronic excessive nutrition also suppresses thermogenesis. Indicating a common mechanism, these cold-induced energy conservation pathways are also paradoxically activated in the setting of obesity^17–20^, which inhibit thermogenic clearance of excess energy, further contributing to obesity and secondary metabolic complications^21–23^. Therefore, the identification of energy conservation pathways that limit thermogenesis should provide strategies for pharmacotherapy against obesity and metabolic comorbidities.

Thermogenesis is initiated at the innermembrane (IM) of BAT mitochondria, where the proton gradient established by the breakdown of lipids and glucose rapidly flows back into the matrix via cold-induced activation of uncoupling protein 1 (UCP1)^5,24^ and/or ATP synthase^25,26^. Ensuing proton accumulation in the matrix removes the bottleneck for the reduction of NAD^+^ and FAD^+^ into NADH and FADH_2_, respectively, which in turn leads to the amplification of the preceding exothermic reactions including the citric acid cycle (TCA)^13^. However, chronic stress from nutrient overload or prolonged thermogenesis continuously reduces IM potential (Δ*Ψ*) due to constant proton leak from the intermembrane space into the matrix^27^. Proton leak-induced drop in local pH was shown to weaken protein-protein interactions due to histidine protonation, which represents the only residue that alters the net charge of a protein upon a small drop in neutral pH (pKA 6.04)^28^: For instance, a matrix soluble protein, Sirtuin 3 (SIRT3), binds ATP synthase and resides at IM at neutral pH. However, proton leak-induced protonation of a histidine on ATP synthase leads to the dissociation and translocation of SIRT3 into the matrix where it alters mitochondrial function by deacetylating a new set of proteins^28^. Based on this premise, we postulated that chronic thermogenic and metabolic uncoupling would cause a sustained drop in pH and increase histidine protonation at the IM-matrix interface. We identified a subset of proteins that translocate from IM into matrix in response to extended cold and caloric stress using proteomic analysis of BAT submitochondrial fractions. Ultimately, we identified a novel mechanism whereby the electron transport chain (ETC) and the mitochondrial calcium uniporter complex (MCUC) serve as docking ports for a subset of soluble matrix proteins. While most translocating proteins supported continued catabolism, a small subset belonged to acyl-CoA thioesterases which prevent the catabolic consumption of stored energy. We tested this new mitochondrial paradigm on acyl-CoA thioesterase 9 (ACOT9), which represented the strongest association to obesity in human subcutaneous white adipose tissue (scWAT) among translocating proteins. Chronic stress-induced matrix localization of ACOT9 blocked excess energy utilization in thermogenesis and promoted obesity and metabolic disorders in mice.

## Results

To promote mitochondrial stress in BAT, we subjected mice to caloric stress by feeding a high fat diet (HFD, 60% fat) *ad libitum* for 10 w, as opposed to regular chow diet (chow, 13% fat) as control. After 10 w, we applied chronic cold stress by housing at 4°C for 2 w, or thermoneutral housing at 30°C for control, which prevents thermogenesis and minimizes proton leak (Fig. 1a). BAT mitochondria were separated into matrix and IM fractions (Extended Data Fig. 1a), and the abundance of proteins in each fraction was measured using mass spectrometry (Supplementary File 1). Overall, 584 proteins were detected in all fractions and conditions, and 375 proteins were detected only in stress conditions (Extended Data Fig. 1b). Among these, 151 were unique to Chow 4°C condition whereas 107 were shared among cold exposure groups regardless of diet, and 102 were shared by all stress conditions compared to Chow 30°C controls. Unsupervised hierarchical clustering and correlation analysis of M/IM ratios indicated reproducibility between biological replicates (Figs. 1b, Extended Data Fig. 1c, and Supplementary Data File 2). Interestingly, the matrix-enrichment of proteins increased with increasing stress conditions (Fig.1c). We confirmed that known IM and matrix markers did not change localization and remained enriched in IM or matrix regardless of testing condition (Extended Data Figs. 1d and e, respectively). Nutritional, extended cold, and both stresses combined resulted in the matrix-enrichment of 55, 71, and 87 IM proteins respectively (Extended Data Fig. 2a). Conversely, using the inverse criteria for identifying a switch in enrichment, we only found 9, 7, and 5 proteins switching from M to IM enrichment, in cold, HFD, or both stresses combined, respectively. Because both chronic nutritional and thermogenic stress exacerbate proton leak into mitochondrial matrix^27,29^, we overlapped all three stress conditions to determine whether reduced Δ*Ψ* via different types of stress would trigger the same factors to translocate from IM into matrix. In a direction that is anticipated from a SIRT3-like mechanism, 73 IM-enriched proteins robustly switched to matrix-enrichment under all stress conditions as opposed to only 4 proteins switching from M to IM (Figs. 1d, e, and Extended Data Figs. 2b and c). To validate human relevance, we monitored breast scWAT from lean and obese women^30^ because, unlike mice, human thermogenic adipocytes are scattered among scWAT^31^. These tissues expressed UCP1 along with several other brown or beige adipocyte markers, 5 of which were inversely associated with BMI^32,33^ (Extended Data Figs. 3a and b). We found that 24 translocating proteins correlated with BMI (Figs. 1f, g, and Supplementary Data File 3).

**Fig. 1.**
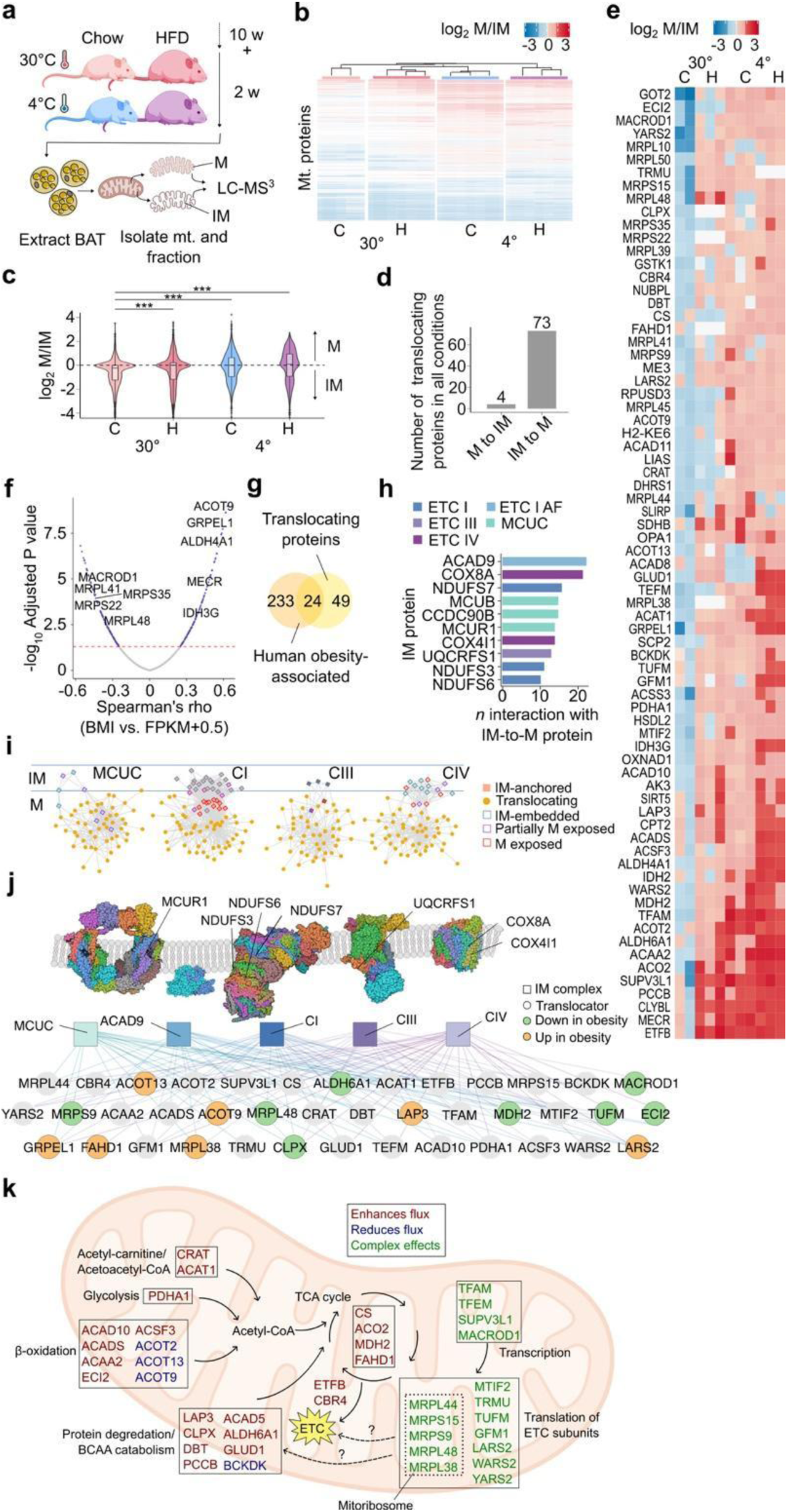
TCA and ETC regulatory proteins dock at the IM-matrix interface in health but translocate to the matrix in stress. **a**, Experimental design. **b**, Unsupervised hierarchical clustering of matrix (M)-to-innermembrane (IM) ratios of detected mitochondrial proteins. **c**, Global shifts in M-IM ratio in response to stress. ***P value < 0.001, one-way ANOVA. **d,** Bar plot showing number of proteins which fit the criteria for either M-to-IM switching or IM-to-M switching. **e**, Subset of proteins which robustly shift from IM to M (“translocators”). **f**, Spearman correlation analysis of 100 scWAT transcriptomes with patient BMI. Genes with significant correlations that were also determined to be shifting IM to M in all stresses are labeled. **g**, Set analysis of human obesity-associated genes and translocators. **h**, Network analysis of known protein-protein interactions between IM-anchored proteins and translocators. **i**, Matrix exposure of subunits with a high degree of connectivity to translocating proteins. **j**, Schematic of IM-anchor to translocator connectivity. Subunits with a high degree of connectivity to translocators are labeled. Individual subunits are collapsed into complex-level nodes. Each edge indicates an experimentally determined protein-protein interaction between subunit and translocator. **k**, Most translocating proteins are enzymes which positively (maroon) affect flux through TCA/ETC; however, several of the enzymes negatively affect flux (blue). Translocation of mitoribosome subunits and other ETC translational machinery has complex effects on flux (green).

Because mitochondrial stress increases the acidity of the matrix^28,34–36^, we next investigated whether the translocation of these 73 proteins were rooted in pH-dependent interactions like SIRT3 and ATP synthase. A mitochondrial interactome analysis between the 73 translocators and IM-embedded proteins (Supplementary Data Files 4 and 5) identified 40 translocators that had experimentally established interactions with only the matrix-exposed subunits of IM complexes. Among these, ETC complexes I (NDUFS3, NDUFS6, and NDUFS7), III (UQCRFS1), and IV (COX8A and COX4I1), ACAD9 (Complex I assembly factor), and three paralogous MCUC subunits (MCUR1, MCUB, and CCDC90B) were disproportionately connected to several translocating proteins (Figs. h-j, Extended Data Fig. 4, and Supplementary Data File 6). Translocating proteins broadly fit into two functional categories: (1) proteins involved in the transcription and mitoribosomal translation of ETC components and (2) enzymes which regulate flux through TCA cycle/ETC (Fig.1k). The translocation of the mitoribosome components correlated with reductions in Complex I and IV subunits but increases in Complex III subunits with convoluted consequences on mitochondrial function (Fig.1k, green and Extended Data Figs. 5a and b). On the other hand, there was a clear direction toward catabolic activation by the translocation of many TCA cycle/ETC components (Fig.1k, orange). Nonetheless, four enzymes that limit acyl-CoA and branch-chain amino acid (BCAA) catabolism also translocated to potentially taper excess energy loss to thermogenesis (Fig.1k, blue).

Because each identified IM anchor was disproportionately associated with multiple translocating proteins (Fig.1j), we next investigated the prospects of a common docking site at each IM anchor. Histidine is the only amino acid that is subject to protonation at physiologically relevant pH fluctuations such as proton leak-induced drop in pH from 7.4 to 6.0 at the matrix interface of the IM^28,33–36^. Persistent histidine protonation can disrupt cation-π and π-π stacking interactions^37^ and weaken soluble protein-docking at the IM as in the case of SIRT3 dissociation from ATP synthase^28^. We identified histidine-dependent cation-π and π-π interactions on the most highly connected IM proteins and translocators. Screening 1,175 AlphaFold^38^ multimer models yielded 201 predicted histidine-dependent cation-π and π-π interactions (Fig. 2a and Supplementary Data File 7). NDUFS6, NDUFS7, ACAD9, UQCRFS1, and COX8A, were disproportionately well connected to translocators via histidine-dependent interactions (151/201 predicted interactions). By algorithmically embedding solved complex models into models of IM bilayers, we determined the feasibility of these histidine-dependent interactions, given exposure to the matrix (Fig. 2b and Extended Data Fig. 5). Overall, our findings support that IM-anchored complex subunits have reversible, pH-sensitive cation-π and π-π docking interactions with dozens of TCA/ETC regulators. Moreover, we find these interaction surfaces are biased towards histidine- or histidine acceptor-rich hotspots^39^ on IM proteins (Figs. 2c-g).

**Fig. 2.**
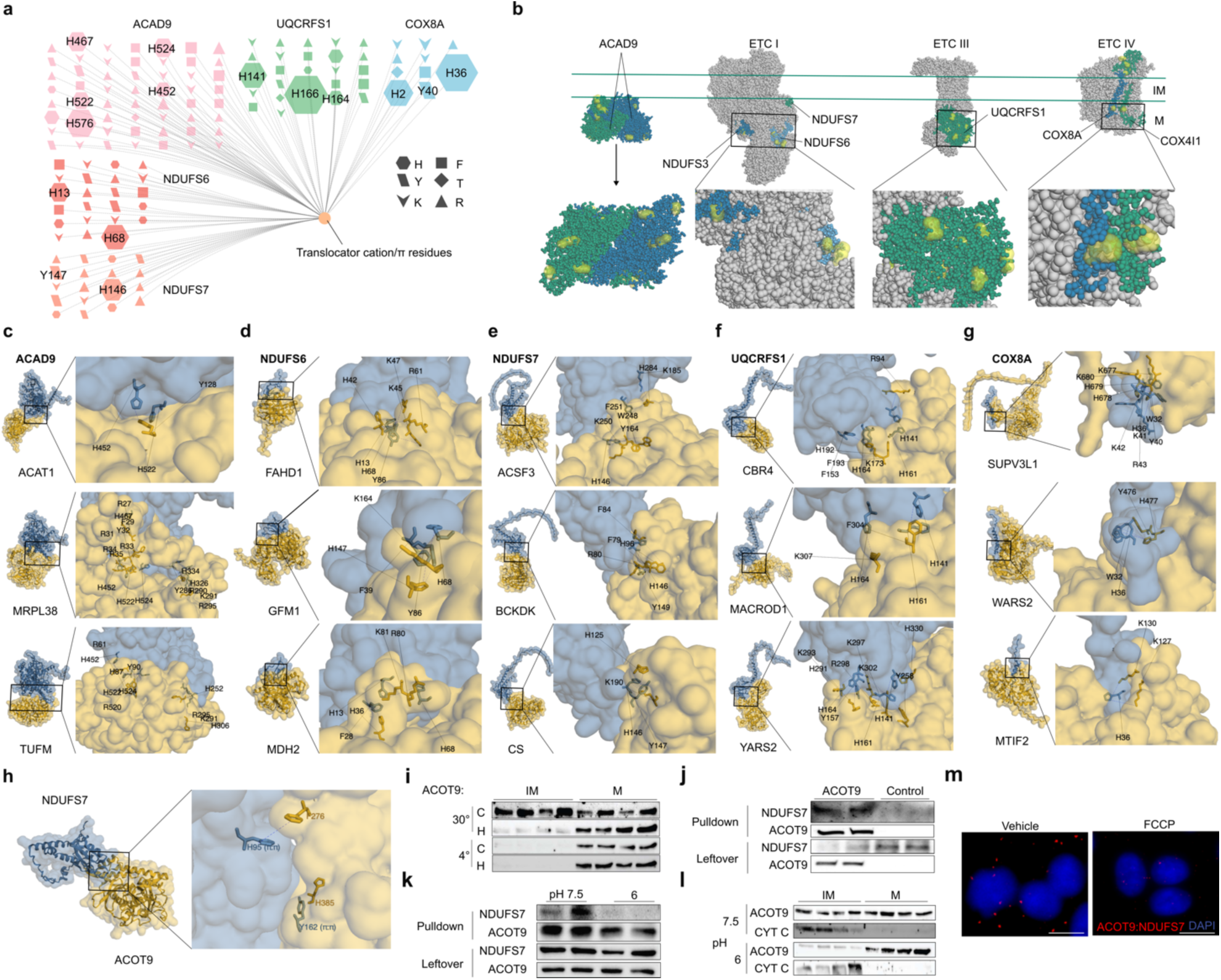
pH-sensitive interactions between IM and translocating proteins. **a**, Graph analysis of predicted IM subunit-to-translocator cation-π and π-π interactions that involve histidine residues. **b**, Matrix exposure of histidine-rich hotspots on IM-anchored complexes. Models of translocator (yellow) and IM-anchored protein (blue) multimers for: **c**, ACAD9, **d**, NDUFS6, **e**, NDUFS7, **f**, UQCRFS1, and **g**, COX8A. In **c** to **h**, inset shows possible cation-his and pi-his interactions. **h**, Model of human IM-anchored subunit NDUFS7 (ETC I) docking with translocator ACOT9. **i**, ACOT9 expression in the IM and matrix fractions of BAT mitochondria from mice fed chow (C) or high-fat diet (H) in thermoneutrality (30°) or cold (4°). **j**, Recombinant ACOT9 fused to maltose binding protein (MBP-ACOT9) or MBP alone as negative control were used to pulldown NDUFS7 from HEK293E protein lysate. **k** ACOT9-NDUFS7 co-immuno-precipitation at pH 7.5 vs. 6.0. **l**, ACOT9 immunoblot from fractionated BAT mitochondria at pH 7.5 vs. 6.0. **m**, *In situ* proximity ligation assay (PLA) in HEK293 cells where Texas Red puncta indicate ACOT9-NDUFS7 interactions in control (Vehicle (Veh.) only) conditions and after induction of proton leak by ionophore carbonyl cyanide-*p*-trifluoromethoxy phenylhydrazone (FCCP).

Our primary goal was to uncover a new mechanism that prevents excess energy loss; thus, we trained our focus on 4 proteins with negative feedback on thermogenesis (Fig. 1k, blue). Interestingly, most of these proteins belonged to the ACOT family (ACOT2, ACOT9 and ACOT13), which have been previously associated with thermogenesis^40,41^. Among these proteins, the enzymatic activity of ACOT9 is the common checkpoint for mitochondrial catabolic reactions as it hydrolyzes and deactivates acetyl-CoA^42^ before its entry into the TCA cycle^43^. ACOT9 also represented the tightest correlation with increased BMI in humans among all translocating proteins (Spearman’s rho=0.62). We have previously shown that mice with whole body ablation of ACOT9 (*Acot9^−/−^*) exhibit increased energy expenditure (EE) and protection against HFD-induced weight gain and adiposity^40^, but the mechanism remained unexplored. Therefore, we expanded testing of this new concept of submitochondrial translocation in thermogenic regulation to ACOT9 as a model mechanism.

To further address whether the translocation of ACOT9 from IM into matrix followed a pH-dependent mechanism as seen with SIRT3 and ATP synthase^28^, we tested the pH dependence of the interactions between ACOT9 and its interacting partner at the IM, the ETC Complex I subunit NDUFS7^44^. *In silico* prediction model for the ACOT9-NDUFS7 complex revealed two histidine residues (ACOT9 His385 and NDUFS7 His95) on the interaction surface (Fig. 2h). We first confirmed that ACOT9 was more abundant in the IM than the matrix of BAT mitochondria in the absence of any stress but increasingly translocated into the matrix upon caloric stress, chronic cold, and both stresses combined in immunoblots (Fig. 2i). In pulldown assays, ACOT9 interacted with NDUFS7 at resting mitochondrial pH of 7.4, underlying its localization to the IM in the absence of mitochondrial stress (Fig. 2j). We next tested the impact of low pH on ACOT9 interactions with NDUFS7 and found that ACOT9 failed to pulldown NDUFS7 when the pH of the assay buffer was lowered from 7.4 to 6.0 to simulate proton leak in low Δ*Ψ* (Fig. 2k). In addition, fractionation of submitochondrial compartments using a low pH buffer resulted in the accumulation of most ACOT9 protein within the matrix compartment even in the absence of any preceding stress, while another peripheral soluble protein, cytochrome C, persistently remained at the IM compartment as control (Fig. 2l). The influence of pH on ACOT9-NDUFS7 interactions was also confirmed by proximity ligation in live cells, which was disrupted upon promoting proton leak with an ionophore (FCCP) (Figs. 2m). Taken together, these findings demonstrate a pH-dependent mechanism by which mitochondrial factors such as ACOT9 can be regulated.

To determine whether a translocator such as ACOT9 could affect thermogenesis in mice, we ramped down the ambient temperature from thermoneutrality to room temperature and then to cold, which gradually increased the biological demand for thermogenesis (Extended Data Fig. 7). Indirect calorimetry measurements indicated that the maximal oxygen consumption (VO_2_) and EE of *Acot9^−/−^* mice diverged from wildtype (*WT*) littermates (*Acot9^+/+^*) as the demand for thermogenesis increased. Suggestive of impaired initiation of torpor, *Acot9^−/−^* mice maintained their core body temperature even after 2 w of mild cold exposure (10°C) unlike *WT* counterparts (Extended Data Fig. 8). To determine whether increased EE in *Acot9^−/−^*mice was due to BAT activity and not other futile mechanisms^45^, thermogenesis was specifically activated by injecting mice with the β_3_ adrenergic receptor (β_3_-AR) agonist CL-316,243. We found that in *Acot9^−/−^* mice, VO_2_ rates were markedly higher than *WT,* regardless of the diet upon BAT activation (Fig. 3a). On the other hand, physical activity and digestive efficacy did not contribute to the altered energy homeostasis between *Acot9^−/−^* and *WT* mice (Extended Data Figs. 9a-c). Because BAT is the main hub for thermogenesis, we studied the impact of ACOT9 in BAT by generating mice with BAT-specific ablation of ACOT9 (*Acot9^B-KO^*) (Extended Data Figs. 10a and b). In a trend similar to *Acot9^−/−^* mice^40^, *Acot9^B-KO^* mice gained 13% less body weight compared to *Acot9^B-WT^* littermates after 10 w on HFD (Fig. 3b), which was attributable to 43% weight reduction in total fat mass (Fig. 3c). Unlike fat mass, lean mass remained unchanged in these mouse lines (Extended Data Figs. 10c-f). Despite reduced adiposity, *Acot9^B-KO^* mice consumed 10% more food than *Acot9^B-WT^* under thermogenic conditions (Fig. 3d). The compensatory aspect of increased food intake was supported by *Acot9^B-KO^*mice losing 30% more weight than *Acot9^B-WT^* following a 6-h fast when they were unable to make up for increased thermogenic energy expenditure by eating more (Fig. 3e). Indirect calorimetry analysis of *Acot9^B-KO^* mice indicated that the loss of ACOT9 expression in BAT alone was sufficient to increase VO_2_, VCO_2_, and EE compared to *Acot9^B-WT^* counterparts (Fig. 3f and Extended Data Figs. 11a and b). During the indirect calorimetry analysis, food intake remained higher for *Acot9^B-KO^* mice compared to *Acot9^B-WT^* littermates (Fig. 3g), but there was no significant difference in physical activity (Extended Data Fig. 11c).

**Fig. 3.**
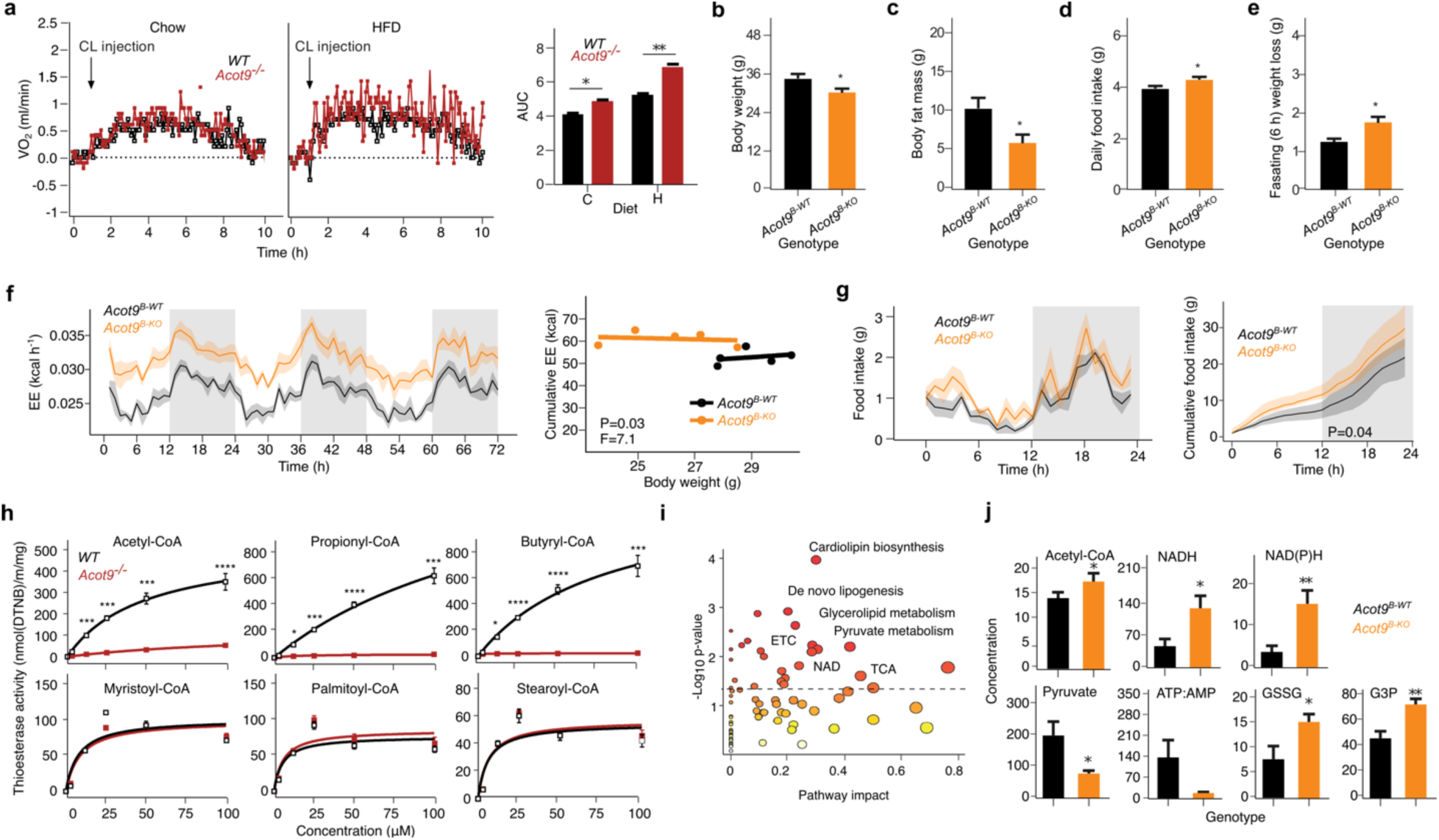
The loss of ACOT9 protects against adiposity by promoting thermogenesis. **a**, Oxygen consumption rates (VO_2_) and area under the VO_2_ curve values of HFD and chow-fed *Acot9^−/−^* and WT mice (n=6) in response to thermogenic stimulation by CL-316,243 injection (1 mg/kg body mass). *P<0.05, ***P<0.001, Student’s t-test. (**b** to **j**) Metabolic data from 10 w HFD-fed *Acot9^B-KO^* and *Acot9^B-WT^* mice (n = 8 / genotype). **b**, Total body weight. **c**, EchoMRI measurement of total fat mass. **d**, Daily food intake under thermogenic condition (4°C). **e**, Weight loss following 6-h fast. **f**, Hourly and cumulative energy expenditure (EE; P < 0.05 F = 7.1, analysis of covariance). **g**, Hourly and cumulative food intake (kcal) over 3 d period. **h**, Thioesterase activity in the BAT of HFD-fed *Acot9^−/−^* and WT mice against short- and long-chain fatty acids. *P<0.05, ***P<0.001, ****P<0.0001, Michaelis-Menten. **i**, Metabolic pathway analysis of HFD-fed *Acot9^B-KO^* vs. *Acot9^B-WT^* mouse BAT was determined by mass spectrometry-based metabolomics. **j**, Metabolites that are directly implicated in mitochondrial activity in the BAT of HFD-fed mice. In bar graphs, bars indicate the mean and error bars indicate ± SEM, *P<0.05, **P<0.01, independent samples t-test.

To elucidate the significance of ACOT9’s enzymatic activity in BAT, we first confirmed that the deletion of ACOT9 in BAT did not alter the expression of other ACOT family members (Extended Data Fig. 12). We next verified that the thioesterase activity of ACOT9 in BAT is specifically directed toward the hydrolysis of short-chain acyl-CoA such as acetyl-CoA (2C), propionyl-CoA (3C), and butyryl-CoA (4C), but not mid- or long-chain acyl-CoA species (Fig. 3h). Considering that propionyl-CoA is the breakdown end-product of odd-chain acyl-CoA, which represent less than 1% of acyl-CoA^46^ and that butyryl-CoA is further oxidized into acetyl-CoA, these results solidify the role of ACOT9 as a key checkpoint for acetyl-CoA flux into TCA cycle. Mass spectrometry-based metabolomics analysis of BAT further supported the role of ACOT9 in limiting the utilization of acetyl-CoA in pathways essential to thermogenesis, including TCA cycle/ETC, pyruvate and butyrate metabolism, nicotinate and nicotinamide metabolism, fatty acid (FA) oxidation, glycerolipid metabolism, *de novo* lipogenesis, and cardiolipin biosynthesis (Fig 3i, Supplementary Data File 8). Among BAT metabolites, acetyl-CoA, NADH, NADPH, glutathione/oxidized GSH (GSSG), and glycerol-3-phosphate abundance trended higher, whereas ATP: AMP and pyruvate levels were lower in *Acot9^B-KO^* mouse BATs compared to *Acot9^B-WT^* counterparts (Fig. 3j).

Because thermogenesis is primarily fueled by β-oxidation following lipolysis, we assessed basal and isoproterenol-stimulated lipolysis rates in *Acot9^−/−^* and *WT* primary BAT cultures. In further support of improved FA utilization and disposal, the loss of ACOT9 increased overall lipolysis levels (Fig. 4a). Nonetheless, increased thermogenesis was not additionally supplemented by enhanced WAT lipolysis as evidenced by a lack of difference in *ex situ* lipolysis of WAT (Extended Data Figs. 13a and b). ACOT9 also inhibited β-oxidation as evidenced by decreased basal and norepinephrine (NE)-stimulated oxygen consumption rates (Fig. 4b), as well as breakdown of radiolabeled (^14^C)-palmitate in complementary assays (Fig. 4c). Although the rates of FA uptake, *de novo* lipogenesis, and overall lipid accumulation remained unchanged (Extended Data Figs. 14a-d), mRNA abundance for fatty acid translocase (CD36) and carnitine palmitoyltransferase 1 (CPT1) was higher in *Acot9^B-KO^* mice BATs vs. *Acot9^B-WT^* counterparts (Extended Data Fig. 15a). Lipidomics analysis indicated that ACOT9 promoted the accumulation of saturated triglyceride species (Extended Data Figs. 15b and c, and Supplementary Data File 9). Overall, these findings emphasize the significance of ACOT9 as a key checkpoint in FA utilization in BAT.

**Fig. 4.**
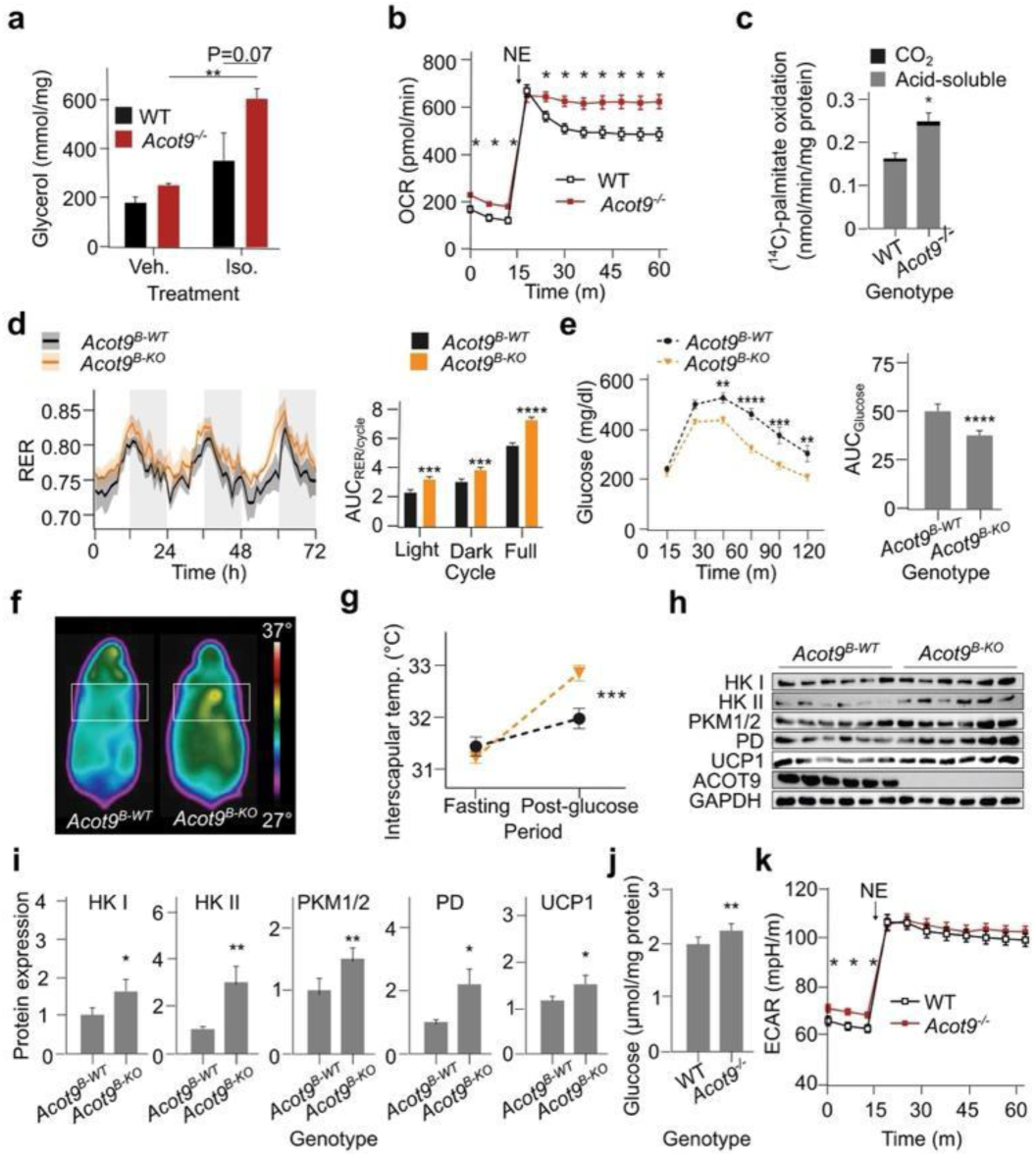
BAT-specific deletion of ACOT9 improves excess energy utilization via thermogenesis. (**a** to **c**) Data from cultured *Acot9^−/−^* and WT primary brown adipocytes. **a**, Lipolysis was determined as a function of released glycerol following stimulation with 1 µM isoproterenol or dimethyl sulfoxide (DMSO) for vehicle. **b**, Basal and norepinephrine-stimulated (5 µM final concentration/well) oxygen consumption rates (OCR) as determined by seahorse flux analysis. **c**, Fatty acid oxidation as determined by the breakdown of ^14^C-labelled palmitic acid into CO_2_ and acid-soluble metabolites. **d**, Respiratory exchange ratio (RER) of HFD-fed *Acot9^B-KO^*and *Acot9^B-WT^* mice at thermogenic condition (4°C) (n=8). Bar graphs reflect areas under the RER curves (AUC) during light, dark, and full day. **e**, Systemic glucose tolerance in HFD-fed *Acot9^B-KO^*and *Acot9^B-WT^* mice tested by intraperitoneal glucose tolerance test (IpGTT; 2 g/kg body mass) (n=8). Bar graph represents AUC. **f** and **g**, Thermographic analyses of HFD-fed *Acot9^B-KO^* and *Acot9^B-WT^* mice dorsal area measured at fasting and 30 m after glucose injection (2 g/kg body mass) (n=6). Analyzed by linear regression t-test on the slope (***P<0.001). **h**, Immunoblot showing relative abundance of proteins that are involved in glycolysis (hexokinase I (HK I), HK II, pyruvate kinase 1/2 (PKM1/2), and pyruvate dehydrogenase (PD)) and thermogenesis (uncoupling protein 1 (UCP1)) in BAT of HFD-fed *Acot9^B-KO^* and *Acot9^B-WT^* mice was determined by immunoblot and **i**, densitometric analysis. **j**, Glucose uptake in cultured *Acot9^−/−^* and WT primary brown adipocytes. **k**, Basal and norepinephrine-stimulated (5 µM final concentration/well) extracellular acidification rates (ECAR) of cultured *Acot9^−/−^* and WT primary brown adipocytes. Bar plots show mean ± SEM. *P<0.05, **P<0.01, ***P<0.001, and ****P<0.0001, independent samples t-test.

β-oxidation of lipids is not the sole type of fuel for thermogenesis. An increasing number of studies have established glucose as a substantial cellular fuel that feeds thermogenesis through intermediary mechanisms in BAT^9,47–49^. Noticeably, both HFD-fed *Acot9^B-KO^* (Fig. 4d) and *Acot9^−/−^*(Extended Data Fig. 16) mice exhibited higher utilization of carbohydrates as fuel, which was evidenced by increased respiratory exchange ratio (RER). We therefore assessed glucose tolerance in weight-matched *Acot9^B-KO^* mice and *Acot9^B-WT^* littermates (Fig. 4e). BAT-specific ablation of ACOT9 resulted in 25% faster glucose disposal in HFD-fed mice without altering fasting blood glucose. To determine whether this difference was due to glucose clearance via BAT thermogenic activity, we tested glucose-induced thermogenesis using intrascapular thermal imaging^50^. A bolus dose of glucose injection increased heat dissipation in dorsal area of *Acot9^B-KO^* mice by 0.9°C higher than *Acot9^B-WT^* littermates (Figs. 4f and g), while core body temperature remained stable (Extended Data Fig. 17). In line with these findings, the deletion of ACOT9 accelerated BAT glycolytic metabolism as evidenced by upregulation of key glycolysis enzymes in *Acot9^B-KO^*mice BATs vs. *Acot9^B-WT^* counterparts (Figs. 4h and i). Likewise, UCP1 expression was also elevated in BAT of *Acot9^B-KO^* mice. In further support of improved glucose utilization, we found that primary brown adipocytes lacking ACOT9 exhibited significant increases in both glucose uptake and extracellular acidification rate (ECAR) despite the saturated insulin concentration in the primary brown adipocyte maintenance media (Figs. 4j and k). These data support the notion that BAT cells upregulate pathways to break down glucose into acetyl-CoA and dissipate it as heat in the absence of an active ACOT9 within the matrix.

## Discussion

Recent studies on mitochondrial structural and functional proteomics have outlined how remarkably complex its protein landscape is^51–53^. Δ*Ψ* is a critical component of mitochondrial localization in presequence-containing proteins via the TIM23 complex^54–56^. In addition, cold-induced mitochondrial protein import is essential for mitochondrial integrity^57^. Our results expand a new concept of Δ*Ψ*-mediated protein translocation from IM surface to Matrix, first described in SIRT3^28^, to over 70 additional proteins that also display dynamic submitochondrial localization in response to altered Δ*Ψ*. The most important finding in the present study is that excessive proton accumulation at the matrix interface of IM has the potential to weaken protein-protein interactions and cause translocation of an IM-bound protein into the matrix. The data thus introduce a new paradigm that concerns overall metabolic health through the scope of a new mitochondrial stress response.

The interactome analyses revealed common histidine-dependent binding pockets whereby protonation can promote the disassociation of proteins from the IM by two primary mechanisms, the first of which is by switching the attraction between the two proteins into repulsion as in the case for ACOT9 and NDUFS7. Secondly, histidine residues on a protein would translate to a higher net charge at low pH, which can alter protein-folding and stability^18,58,59^. While we tested this model in ACOT9 and NDUFS7, the protonation state of key histidines under low pH remains to be identified, and the nature of interactions for each interacting partner will have to be elucidated one at a time based on the unique structure of each protein. In addition to disrupting the interactions between an IM anchor and its matrix-translocating partners, histidine protonation may also impact the interactions among the IM anchors as ETC complexes I, III and IV form the respirasome supercomplex to enhance ETC activity. Although we have identified these exact respirasome components to be the most highly connected IM anchors, additional studies will be required to determine whether the formation of the respirasome is inhibited by the binding of the translocating proteins to the respirasome components, whereby their dissociation would in turn promote the formation of the respirasome supercomplex to enhance ETC activity.

Our data suggest that the submitochondrial localization of an enzyme might be important for two main reasons: (1) the enzyme might be dormant or active while docked at the IM, or (2) the enzyme might be restrained and away from its specific substrate(s). Our extension of this concept to ACOT9 indicates that ACOT9 lays dormant while in interaction with NDUFS7 as monomeric ACOTs are catalytically inactive^42,60^. Thus, its dissociation from NDUFS7 would not only give way to its activation by multimerization but also expose it to its substrate acetyl-CoA within the matrix. Reciprocally, the binding of ACOT9 to NDUFS7 may have improved complex I activity, making its stress-induced matrix localization an unfavorable condition. However, this would not explain the improved metabolism of mice lacking ACOT9 expression. Nonetheless, the potential impact of binding by a matrix protein on the activity of an IM complex should not be ignored moving forward. Likely evolved and conserved to prevent uncontrolled energy loss via thermogenesis, our results indicate that ACOT9 promotes obesity and related metabolic disorders given the current epidemic of excessive calorie consumption. Targeted inhibition of ACOT9 might increase energy expenditure and thereby be useful in the treatment of metabolic disorders.

In the current study, we specifically investigated conditions that would cause chronic and steady mitochondrial stress under extended cold and high fat diet, which would heavily tilt the equilibrium of submitochondrial protein localization and facilitate the identification of translocating proteins. Nonetheless, it is possible that even short-term physiological changes in Δ*Ψ*, such as short bouts of fasting, food intake, cold exposure, daily torpor and glucocorticoid signaling could cause IM-to-M translocation to regulate mitochondrial activity. After all, healthy biological functions such as ATP synthase activity alone can induce a temporary drop in local pH, and thermogenesis consistently operates at low Δ*Ψ* whether it is acute or chronic. This notion for acute control of thermogenesis for energy conservation is further supported by other checks and balances including by ACOT11 and AC3-AT^18,57^. Therefore, the lack of acute stress conditions in the current study is a shortcoming that prevents distinguishing the beneficial IM-to-M translocation events during healthy stress oscillations from those that are maladaptively exacerbated under chronic stress. Nonetheless, even under chronic stress conditions, the majority of translocating proteins consisted of supporters of thermogenesis while only a small subset worked against it (Fig. 1k, 21 orange vs. 4 blue). Finally, although we took advantage of BAT’s unique physiology to observe maximal proton surge from the intermembrane space into the matrix, mitochondrial stress and dysfunction are detrimental for a wider array of disorders including but not limited to developmental, neurological, cardiovascular, and metabolic problems as well as immunodeficiency and cancer^42,61^. In this context, IM-to-M protein translocation does not only represent another rheostat for uncontrolled thermogenesis but also carries the potential to have broader implications in health.

## Materials and Methods

### Animal care

Experimental procedures were approved by and performed in compliance with the Institutional Animal Care and Use Committee protocol at the Weill Medical College of Cornell University (New York, NY). Husbandry and housing were conducted at the Weill Cornell Medicine Animal Facility under strict biosecurity measures. Unless indicated, mice were housed in a clean room set at 22°C (room temperature) with regulated humidity on a daily 12-hour light/dark cycle with access to drinking water and feeding diet. RD caloric intake (kcal) was comprised of proteins (24%), fat (13%), and carbohydrates (62%) (LabDiet PicoLab Rodent Diet 20 [5053]; St. Louis, MO, USA). For caloric stress, overnutrition was instated by feeding mice a HFD with 60% kcal from fat (Research Diets Rodent Diet [D12492]; New Brunswick, NJ, USA). This study did not include time-restricted dietary feeding; all diets were fed *ad libitum* after weaning at 4 w of age for the specified periods.

### Acclimation to ambient temperature

All mice were maintained at 22°C until they are assigned for acclimation for specific experiments. Thermoneutrality-based experiments were conducted by housing mice at 30°C at the Metabolic Phenotyping Center of Weill Cornell Medicine. Accordingly, mice were housed in an environmental enclosure (Darwin Chambers Laboratory Incubator [DB034]; St. Louis, MO, USA) at 30°C for 2 w before proceeding with tissue dissection. For cold acclimation, mice were transferred to the temperature- and humidity-controlled cold room (4°C) at the Research Animal Resource Center at Weill Cornell Medicine. For mice to tolerate an abrupt change in ambient temperature from 22°C to 4°C, all mice were kept in the original cages with bedding and sterilized igloos (Bio-Serv [K3352]; Flemington, NJ, USA) for 3 d in the cold room. After that, mice were split in pairs, and igloos and 70% of bedding were removed from the cages for another 4 d. Following this period, mice were single housed for 2 w in the cold room with no igloos and minimal bedding (<10%) in the cages. Food intake during the cold exposure was measured with a lab-designed in-cage metal feeding hopper for the accurate measure and prevention of feed loss into bedding.

### Proteomic sample preparation

Following the protein quantification with Pierce Bicinchoninic Acid (BCA) Protein Assay Kit – Reducing Agent Compatible (ThermoFisher Scientific [23250]; Waltham, MA, USA), 5 μg of each mitochondrial fraction protein lysate was moved into a clean 1.5-ml tube. After setting up the tubes ready, each tube was brought to a final volume of 300 μl by addition of phosphate-buffered saline (PBS), followed by precipitation with trichloroacetic acid (MilliporeSigma [T6399]; St. Louis, MO, USA) to a final concentration of 25%, vigorously vortexed and incubated on ice overnight. Trichloroacetic acid precipitates were centrifuged at 21,130 x g for 30 m at 4°C, washed twice in 500 µl of ice-cold acetone, and centrifuged at 21,130 x g for 10 m after each wash. Following precipitation and washes, pellets were allowed to completely dry at room temperature. Dry pellets were resuspended in 100 μl of 100 mM triethylammonium bicarbonate (TEAB) (ThermoFisher Scientific [90114]; Waltham, MA, USA) and 0.5% sodium dodecyl sulfate (SDS), and reduced with 9.5 mM tris-carboxyethyl phosphine (TCEP) for 60 m at 55°C. Following reduction of disulfide bonds with TCEP, the denatured protein mixture was centrifuged at 21,130 x g for 5 m then alkylated with 4.5 mM iodoacetamide for 30 m in the dark at room temperature. After reduction and alkylation of disulfide bonds, the denatured protein mixture was precipitated out of solution by addition of 600 µl of ice-cold acetone and placed in the -20°C freezer overnight. The following day, proteins were pelleted by centrifugation at 8,000 x g for 10 m and allowed to air-dry at room temperature. Protein pellets were then reconstituted in 100 µl 100 mM TEAB and 1 mM CaCl_2_. Enzymatic digestion was carried out in the dark on a thermal shaker (for 16 h at 37°C) following the addition of 0.1 µg of sequencing grade Trypsin (Promega [V5111]; Madison, WI, USA).

### Tandem mass tag (TMT) labeling

TMT labeling was performed generally as per manufacturer’s protocol. Briefly, each TMT tag was resuspended in 41 μl anhydrous acetonitrile with intermittent vortexing for 10 m. Following resuspension, tags were added to corresponding mitochondrial fractions and labeling reaction was allowed to proceed for 1 h at room temperature. Reactions were quenched by the addition of 8 μl of 5% hydroxylamine in 100 mM TEAB and incubated for 15 m. Labeled mitochondrial fractions were pooled and desalted on 1 cc/50 mg Sep-Pak C18 cartridges (Waters Corporation [WAT054955]; Milford, MA, USA) on a vacuum manifold, and desalted peptides were dried down in a SpeedVac Concentrator (ThermoFisher Scientific; Waltham, MA, USA). Dried peptides were reconstituted in 300 µl of 0.1% trifluoroacetic acid (TFA) in water. Pierce High pH Reverse-Phased Fractionation Kit spin columns ([84868]; ThermoFisher Scientific; Waltham, MA, USA) were equilibrated, and samples were fractionated per manufacturer’s instructions into 8 fractions, 2 washes and a flow-through fraction (11 total) and dried in a SpeedVac Concentrator. Peptide fractions were reconstituted in 12 µl of 5% acetonitrile and 0.1% TFA in water and transferred to virgin polypropylene vials for multi-notch Ultra-High-Performance Liquid Chromatography-Tandem Tribrid Mass Spectrometry (UHPLC-MS^3^) analysis.

### LC-MS^3^ data acquisition

High pH reverse-phased fractions were run on a 4-h instrument method with an effective linear gradient of 180 m from 5% to 25% mobile phase B with the following mobile phases: A) 0.1% formic acid in water, B) 80% acetonitrile/0.1% formic acid in water on a 50 cm Acclaim PepMap Rapid Separation (RSLC) C18 column (ThermoFisher Scientific [164540]; Waltham, MA, USA) operated by a Dionex UltiMate 3000 RSLCnano system (ThermoFisher Scientific; Waltham, MA, USA) with column heating at 50°C connected to an Orbitrap Fusion Lumos MS^3^. Briefly, the instrument method was a data-dependent analysis and cycle time set to 3 s, total. Each cycle consisted of one full-scan mass spectrum (400-1,500 m/z) at a resolution of 120,000, RF Lens: 60%, maximum injection time of 100 millisecond followed by data-dependent MS/MS spectra with precursor selection determined by the following parameters: Automatic gain control (AGC) target of 4.0e^5^, maximum injection time of 100 millisecond, monoisotopic peak determination: peptide, charge state inclusion: 2-7, dynamic exclusion 10 s with an intensity threshold filter: 5.0e^3^. Data-dependent MS/MS spectra were generated by isolating in the quadrupole with an isolation window of 0.4 m/z with collision-induced dissociation (CID) activation and corresponding collision energy of 35%, CID activation time of 10 millisecond, activation Q of 0.25, detector type Ion Trap in Turbo mode, AGC target of 1.0e^4^ and maximum injection time of 120 ms. Data-dependent multi-notch MS^3^ was done in synchronous precursor selection mode (SPS, multi-notch MS^3^) with the following settings: Precursor Selection Range; Mass Range 400-1200, Precursor Ion Exclusion Properties m/z Low: 18 - High: 5, Isobaric Tag Loss Exclusion Properties: TMT. Number of SPS precursors was set to 10 and data-dependent MS^3^ was detected in the Orbitrap (60,000 resolution, scan range 120-500) with an isolation window of 2 m/z higher-energy collisional dissociation (HCD) activation type with collision energy of 55%, AGC target of 1.2e^5^ and a maximum injection time of 150 millisecond. Raw files were parsed into MS1, MS2, and MS3 spectra using RawConverter^62^.

### Primary LC-MS^3^ data analysis

Data generated were searched using the ProLuCID algorithm^36^ in the Integrated Proteomics Pipeline software platform. Mouse proteome data were searched using concatenated target/decoy UniProt databases. Basic searches were performed with the following search parameters: HCD fragmentation method; monoisotopic precursor ions; high resolution mode (3 isotopic peaks); precursor mass range 600-6,000 and initial fragment tolerance at 600 ppm; enzyme cleavage specificity at C-terminal lysine and arginine residues with 3 missed cleavage sites permitted; static modification of +57.02146 on cysteine (carboxyamidomethylation), +229.1629 on N-terminal and lysine for TMT-10-plex tag; 4 total differential modification sites per peptide, including oxidized methionine (+15.9949), and phosphorylation (+79.9663) on serine, threonine, and tyrosine; primary scoring type by X corr and secondary by Z score; minimum peptide length of six residues with a candidate peptide threshold of 500. A minimum of one peptide per protein and half-tryptic peptide specificity were required. Starting statistics were performed with a Δmass cutoff of 10 ppm with modstat and trypstat settings. False-discovery rates of protein (pfp) were set to 1% (for unenriched datasets). TMT quantification was performed using the isobaric labeling 10-plex labeling algorithm, with a mass tolerance of 5.0 ppm or less. Reporter ions 126.127726, 127.124761, 127.131081, 128.128116, 128.134436, 129.131417, 129.13779, 130.134825, 130.141145, and 131.13838 were used for relative quantification.

### Secondary LC-MS^3^ data analysis

Proteins identified by only one peptide were excluded, as were those annotated as “reverse” or “contaminant.” Missing intensity values were imputed using a small constant (634.29). Fractional localization shifts were calculated by determining the ratio of mitochondrial inner membrane (M) to mitochondrial matrix (IM) intensities. These ratios underwent log_2_ transformation for subsequent statistical analyses.

Mitochondrial proteins were annotated according to the MitoCarta3.0 mouse database^63^. Proteins specifically localized to the mitochondrial IM or matrix were selected for further analysis. Proteins were classified based on their localization shifts between the IM and matrix under stress versus control conditions. Proteins classified as translocating from the IM to the matrix had predominantly IM localization in control conditions (log2(M/IM) < 0), exhibited increased matrix localization under stress relative to control (stress log2(M/IM) > control log2(M/IM)), and demonstrated predominantly matrix localization under stress conditions (stress log2(M/IM) > 0.25). Heatmap visualization and unsupervised hierarchical clustering were performed using the pheatmap package.

### Correlation analysis of 100 scWAT sample transcriptomes and patient BMI

Transcriptomic and patient data were acquired from a previously published dataset^30^ and analyzed with consultation from original biostatisticians. Briefly, human subcutaneous adipose tissue was collected at the time of mastectomy following informed consent. The collection of clinical data, e.g., BMI and tissue acquisition for RNAseq was approved by the Institutional Review Boards of Weill Cornell Medicine (IRB 1004010984-01) and Memorial Sloan Kettering Cancer Center (IRB 10-040). RNAseq dataset EGAD00001006397 from the European Genome-Phenome Archive (EGA) was analyzed for BMI-associated gene expression. Thermogenic genes were identified based on known markers of thermogenic function. Mitochondrial gene annotations were derived from the MitoCarta3.0 database^63^. Genes with FPKM + 0.5 > 1 were considered expressed. Correlations between gene expression levels and BMI were calculated using Spearman’s rank correlation coefficient (ρ) to accommodate potential non-linear relationships. P-values were adjusted for multiple testing using the Benjamini-Hochberg procedure to control the false discovery rate. Genes with FDR < 0.05 were considered significantly correlated with BMI.

### Mitochondrial IM-matrix interactome construction and graph analysis

Empirically defined protein-protein interactions were compiled from curated interactions identified using Ingenuity Pathway Analysis tool (Qiagen) or from BioGrid and merged into an undirected graph using igraph (R package). Self-loops and duplicate edges were removed. Node-level degree centralities were computed in R with the igraph package.

### Modeling of IM-anchor complex subunits and translocators

Sequences of the most well-connected IM-anchored subunits were multimerized with translocating proteins using Alphafold2-multimer^38^ and ColabFold^64^. Protein sequences of mouse IM-anchored complex subunits and translocators were retrieved from UniProt. Mitochondrial targeting peptides were predicted using TargetP2.0^65^ and then trimmed accordingly prior to modeling.

### IM-anchor complex subunit and translocator Interface network analysis

For each multimer model, a PDBParser object (Biopython v1.79) was used to create a Python object representing the protein structure. All atoms from each chain were compiled into separate lists. A NeighborSearch object was created for each set of atoms to efficiently query atoms within a specified distance threshold of 5 Å. This threshold was chosen to capture interface interactions that typically fall within hydrogen-bonding or other stabilizing interaction distances.

### Visualization of histidine-dependent cation–π and π–π bonding at multimer interface

Python 3.9, Biopython, and Py3Dmol were used to visualize and highlight cation–π or π–π interactions involving histidine residues in multimers. Residues are screened for any pair where one residue is histidine, and the other is cationic (arginine or lysine) or aromatic (phenylalanine, tyrosine, tryptophan, or histidine) within a 5-Å distance. The final views were rendered with Py3Dmol.

### Prediction Model of human ACOT9-NDUFS7 Interaction

The Alphafold2-multimer prediction process was repeated 10 times for human ACOT9 and NDUFS7, and the predicted structure was selected by the highest iPTM score in the analyses. All amino acids in the interaction interface showed a minimum of 70 model confidence score depending on the global superposition of submolecular atoms. The measure for confidence threshold in AlphaFold is called pLDDT value that is based on the local distance difference test (lDDT-Cα), and any score higher than 70 correspond to an accurate structure and help avoid predicted alignment errors^38^.

### Localization and solvent exposure analysis of key histidine residues

The orientation of IM complexes was determined with PPM 3.0^66^ on the Orientations of proteins in membranes database web server (U. of Michigan). Empirically determined complex models acquired from RCSB PDB database^67^ were computationally embedded in a mitochondrial IM bilayer model, and then IM vs. M localization of key histidine residues was determined with ChimeraX^68^. The py3Dmol library was used to generate interactive three-dimensional representations in which the chains of interest were colored, and solvent-accessible surfaces were rendered around histidine residues to illustrate their exposure to solvent.

### Sample preparation for metabolomics

Metabolites were extracted from 50 mg BAT using 500 µl 80% cold (HPLC grade) methanol for polar metabolite profiling. After homogenization, samples were incubated at -80°C for 4 h followed by a centrifugation at 14,000 x g for 20 m at 4°C. Supernatants were transferred into fresh 1.5 ml tubes and lyophilized without heating. Samples were stored at -80°C until performing the assay.

### Targeted metabolomics analysis

Metabolites in a predefined panel that is comprised of intermediates from central regulative mechanisms such as glycolysis and TCA cycle are analyzed by a hybrid quadrupole Orbitrap-based (Q Exactive) mass spectrometer (ThermoFisher Scientific; Waltham, MA, USA) integrated with a Vanquish UHPLC system (ThermoFisher Scientific; Waltham, MA, USA). The Q Exactive MS was run in polarity-switching mode A PEEK coated HPLC column (SeQuant ZIC-HILIC (5µm, 200Å) 150 x 2.1 mm; MilliporeSigma [150464]; St. Louis, MO, USA) was used for metabolite separation. The flow rate was 150 μl/m. Solutions contained 100% acetonitrile (MilliporeSigma [34851]; St. Louis, MO, USA) for A buffer and 0.1% NH_4_OH/20 mM CH_3_COONH_4_ in water for B buffer. Gradients ran from 85 to 30% A buffer in 20 m followed by a wash with 30% A buffer and reequilibration at 85% A buffer. Finally, metabolites were identified based on exact mass within 5 ppm and standard retention times. The relative metabolite quantitation was performed based on the peak area for each metabolite. All data were analyzed by in-house scripts as previously described^69^. Pathway enrichment analysis was conducted using MetaboAnalyst 5.0^70^ including only the metabolites that were enriched by more than 40% (P < 0.05) in BATs of *Acot9^B-KO^*mice compared to *Acot9^B-WT^*. A limited number of reduced metabolites within BATs of *Acot9^B-KO^* mice were individually presented in bar graphs.

### Sample preparation for lipidomics

Sample lipid extraction was conducted as previously reported^69^. Accordingly, extracts were dried using a refrigerated SpeedVac vacuum concentrator (ThermoFisher Scientific; Waltham, MA, USA) and reconstituted with acetonitrile/isopropanol/water mix (65:30:5) containing stable isotope-labelled internal standards (Avanti Polar Lipids [330707]; Alabaster, AL, USA.

### LC-MS/MS analysis of lipids

For chromatographic separation, samples were loaded in a Vanquish UHPLC system (ThermoFisher Scientific; Waltham, MA, USA) with a Cadenza CD-C18 3 µm ODS column (2.1 mm id x 150 mm) (Imtakt USA; Portland, OP, USA) coupled to a hybrid quadrupole Orbitrap-based (Q Exactive) mass spectrometer (ThermoFisher Scientific; Waltham, MA, USA) via an Ion Max ion source with a HESI II probe (ThermoFisher Scientific; Waltham, MA, USA). The mobile phase was comprised of; a. buffer A - 60% acetonitrile, 40% water, 10 mM ammonium formate with 0.1% formic acid and b. buffer B - 90% isopropanol, 10% acetonitrile, 10 mM ammonium formate with 0.1% formic acid. The LC gradient was as follows: 0–1.5 m in 32% buffer B; 1.5-4 m in 32-45% buffer B; 4-5 m in 45-52% buffer B; 5-8 m in 52-58% buffer B; 8-11 m in 58-66% buffer B; 11-14 m in 66-70% buffer B; 14-18 m in 70-75% buffer B; 21-25 m in isocratic 97% buffer B, 25-25.1 m in 97-32% buffer B; eventually followed by 5 m of reequilibration of the column before the next run. Flow rate was 200 μl/m. For the identification of lipids, a data-dependent MS acquisition method was employed. In this approach, each MS survey scan was followed by up to 10 x MS/MS scans run on the most present ions. Data was collected in positive mode. Electrospray parameters were as follows: spray voltage 3.0 kV, heated capillary temperature 350°C, HESI probe temperature 350°C, sheath gas 35 units; auxiliary gas 10 units. Furthermore, MS scans were as follows: resolution, 70,000 (at m/z 200); automatic gain control target, 3e^6^; maximum injection time, 200 ms; scan range, 250-1800 m/z. For MS/MS scans: resolution, 17,500 (at 200 m/z); automatic gain control target, 1e^5^ ions; maximum injection time, 75 ms; isolation window, 1 m/z; NCE, stepped 20, 30, and 40. Raw experimental data files were processed by using MS-DIAL online tool for identification and relative quantitation of lipids^70^.

### Generation of genetic mouse models

All mice were from C57Bl/6J strain. The transgenic conditional *Acot9^+/+^*mice, which had LoxP sites inserted within the 3^rd^ and 5^th^ introns of *Acot9* was generated from an embryonic stem cell clone (HEPD0691_3_C05 0) as previously described^40^. Specific genotype was confirmed by polymerase chain reaction (PCR) using primers: forward 5’-TCAGCTGATGCCTCATTA CCATGG-3’, and reverse 5’-TGCTGACCCACAATTCTAGTCCCC-3’, which provide PCR product sizes 790 bp and 1000 bp for *WT* and *Acot9^+/+^* alleles, respectively. Germline whole-body ACOT9 knockout (*Acot9^−/−^*) mice were generated by breeding *Acot9^+/+^* mice to CAG-Cre mice, after which offspring mice were crossed with *WT* to obtain CAG-Cre-free generations. For the generation of mice with BAT-specific ablation of ACOT9 (*Acot9^B-KO^*), female *Acot9^+/+^* mice were bred with mice expressing Cre recombinase under the BAT-specific promoter control of the UCP1 (Ucp1-Cre, The Jackson Laboratory #024670; Bar Harbor, ME) that were kindly provided by Dr. Timothy McGraw of Weill Cornell Medicine. Accordingly, *Acot9^+/+^ Ucp1-Cre^+/+^* mice were grouped as the *Acot9^B-KO^* mice, whereas *WT Ucp1-Cre^+/+^* or *Acot9^+/+^ Ucp1-Cre^−/−^*mice served as *Acot9^B-WT^* littermates. Presence of Cre transgene was determined using the PCR primers: forward 5’-CATGCCCAGCTGGAAGTGTTT T -3’ and reverse 5’-CTGGCCTGAAGCCTGTATGTAAAC-3’.

### Metabolic monitoring and body composition analysis

Metabolic activity assessment of mice was analyzed at the Metabolic Phenotyping Center core facility, Weill Cornell Medicine. Total lean and fat mass of mice were analyzed by nuclear magnetic resonance spectroscopy using an EchoMRI 3-in-1 Body Composition Analyzer (Houston, TX, USA). Mouse core body temperature was measured by a rectal probe with or without an intraperitoneal (IP) glucose challenge (2 g/kg body mass) based on the experiment using a digital thermometer (JJS Technical Services [WD-35627-00]; Schaumburg, IL, USA). Respirometry and indirect calorimetry values were measured by a Promethion High Definition Multiplexed Respirometry System for Mice (Sable Systems International; Las Vegas, NV, USA). Mice were acclimated to the metabolic cages for a day and readouts were measured thereafter. For the estimation of EE (kilocalories/time) via indirect calorimetry, rates of oxygen consumption (VO_2_) and carbon dioxide production (VCO_2_) were monitored with a sampling frequency of 1 second. EE was calculated by the Weir Equation (=3.94 x VO_2_ + 1.10 x VCO_2_) and shown as the total kcal/time after the adjustment of covariate effect (ANCOVA). F-value of >3.95 was considered as significant. Normalization for body mass and corrected body mass was calculated by VassarStats online tool (http://vassarstats.net). Respiratory activity was acquired in every 5 m with a dwell time of 30 s/cage. RER was calculated from the VCO_2_:VO_2_ ratio to determine metabolic fuel preference, whereby values closer to 1.0 reflected preference for carbohydrates. RER data was analyzed by two-tailed unpaired (independent samples) t-test because RER values have been reported not to differ by body weight^71^. Dietary intake was determined by gravimetric readouts within the Promethion cages. Mouse physical activity (locomotion) was measured by the beam breaks within a matrix of built-in-cage infrared sensors.

### Pharmacological Activation of β3-Adrenoceptor

Thermogenesis was induced pharmacologically using the β3-AR-specific agonist CL 316,243 (MilliporeSigma [C5976]; St. Louis, MO, USA). 4-week-old *Acot9^+/+^* and *Acot9^−/−^* mice were fed HFD and kept at thermoneutrality (30^°^C) for 4 w. Mice were then single-caged and acclimated to Promethion Cage System for one week prior to injecting IP with saline to establish baseline for each mouse. Mice were injected IP with CL 316,243 (1 mg/kg body mass) and VO_2_ response to both saline and CL 316,243 treatments were determined by metabolic monitoring. Thermogenic response to pharmacological activation was calculated from the difference in the area under the curve (AUC) of CL 316,243 and saline. Final values were compared using an ANCOVA.

### Fecal Calorimetry

Digestive efficacy was analyzed as previously described^72^. Briefly, mice were single housed in clean cages, and fecal samples were collected from the bottom of cages after 24 h. Fecal matter was dehydrated for two days and subjected to bomb calorimetry using a combustion calorimeter (6725 Semimicro, Parr Instrument Company; Moline, IL, USA) to determine caloric density. Total daily caloric output was calculated by multiplying total fecal weight with density.

### Assessment of Glucose Homeostasis

Glucose tolerance test (GTT) was conducted in 12-week-old HFD-fed *Acot9^B-KO^* and *Acot9^B-WT^* mice before the divergence of body weights. Mice were fasted for 6 h prior to injecting IP with glucose (2 g/kg body mass). Blood glucose was measured via tail bleed at 0, 15, 30, 60, 90, and 120-m time points by a glucometer (General Electric [GE100]; Boston, MA, USA). Glucose AUC was calculated from baseline (basal fasting blood glucose) using the trapezoid formula in GraphPad Prism (Dotmatics; Bishop’s Stortford, UK).

### Mouse Interscapular Area Infrared Thermography

A thermal camera system (Teledyne FLIR Systems [T430sc]; Wilsonville, OR, USA) was used to assess mouse interscapular region surface temperature after fasting 12 w-old HFD-fed *Acot9^B-KO^* and *Acot9^B-WT^* mice for 6 h. Each mouse was given isoflurane for 30 s to stabilize the mouse posture while taking thermal images captured at an emissivity setting of 0.95 (JPG format). To determine improved utilization of glucose for EE in BAT, same dorsal area (region of interest; ROI) was photographed for thermogenic activity 30 m after injecting mice IP with glucose (2 g/kg body mass). On the camera, the threshold high and low temperatures were set at 38^°^C and 28^°^C, respectively. Only plastic materials were used during this assay, since metal products may have changed the monitored temperatures. Thermographic data from raw images were acquired based on the protocol previously published^50^.

### Tissue Collection

For tissue and organ harvest, food was removed from cages at 8:00 with free access to water. At 14:00, mice were weighed and anesthetized with isoflurane inhalation. Following tissues were dissected after blood collection by intra-cardiac puncture: BAT, epididymal (visceral) and lumbar (subcutaneous) WAT, and liver. Dissected samples were snap frozen in liquid nitrogen and kept at -80°C until use.

### Protein Isolation from Mitochondrial Innermembrane and Matrix Fractions

Mass spectrometry-based proteomics and immunoblots were carried out on submitochondrial fractions that were harvested from C57Bl/6J WT mice. After weaning at 4 w of age, mice were fed HFD or chow *ad libitum* for 10 w and allocated to the following stress groups: a) RD and thermoneutrality (non-stress), b) HFD and thermoneutrality (only caloric stress), c) RD and cold (only cold stress), and d) HFD and cold (both caloric and cold stress). After 2 w in climate control, mice were euthanized, and BAT were fractionated immediately. Briefly, wet tissues were minced and homogenized in a Potter-Elvehjem tube (12 strokes) in 2 ml cold mitochondria isolation (MTiso) buffer as previously described^40,73^. The samples were loaded onto 5 ml 340 mM cold sucrose gradient and centrifuged at 500 x g for 10 m at 4°C. After gently aspirating the lipid layer over the supernatant, samples were centrifuged at 800 x g for 15 m at 4°C to fully remove unbroken cells and nuclei, after which the supernatants were further centrifuged at 10,000 x g for 10 m at 4°C for pelleting the mitochondria. After washing with ice-cold PBS, the outer membrane of the mitochondria was ruptured by suspending the pellets in 1 ml MTiso buffer with 1 mg/ml digitonin (RPI Research Products International [D43065-0.1]; Mount Prospect, IL, USA), and samples were vortexed for 15 m with a multi-tube vortex (ThermoFisher Scientific [02-215-450]; Waltham, MA, USA). Samples were then centrifuged at 10,000 x g for 10 m at 4°C to pellet the IM and matrix compartments. Supernatant containing the outer membrane plus the intermembrane space was discarded. Pellets were washed with ice-cold PBS, resuspended in 150 µl MTiso buffer, and sonicated (15% 10 s twice/sample) to break open the IM. IM and matrix fractions were then separated by ultracentrifugation (100,000 x g for 10 m at 4°C) with the supernatant containing the matrix and the pellet containing the IM fractions. The supernatant samples were added equal volume of cold 2X radioimmunoprecipitation assay (RIPA) buffer (50 mM Tris-HCl (pH 7.5), 150 mM NaCl, 0.35% NP40, 1 mM EDTA, and 0.5% sodium deoxycholate) and snap frozen. Likewise, IM pellets were resuspended in 1X MTiso 1X RIPA buffer and frozen immediately. All buffers were supplemented with protease inhibitor (Apex Bio [K1010]; Houston, TX, USA) and phosphatase inhibitor (Apex Bio [K1013]; Houston, TX, USA) cocktails.

### Protein Isolation from Tissue or Cell Culture

Prior to immunoblot analyses, a small chunk of frozen BAT sample (∼20 mg) was added cold RIPA buffer and 2 ceramic beads in microcentrifuge tubes with screw caps (OMNI International [19-649]; Kennesaw, GA, USA). For homogenization, the program was set to “3 x 30 s (6 m/s)” in the Bead Ruptor Elite homogenizer (OMNI International [19-042E]; Kennesaw, GA, USA). After homogenization, the supernatant was transferred into a new tube and immediately centrifuged at 16,000 x g for 10 m at 4°C. Clear protein extract (supernatant) was transferred into a new tube and diluted by 3 times with RIPA buffer and aliquoted. For cultured cells, wells were rinsed with ice-cold 1X PBS, cell debris was scraped into RIPA buffer and rotated for 20 m at 4°C. Following centrifugation (16,000 x g for 20 m at 4°C), supernatant containing the protein was snap frozen until protein estimation.

### Protein Quantification and Immunoblot Analysis

For all immunoblot analyses, 3 to 50 µg protein was used for each sample per experiment. Sample protein concentrations were estimated by the Pierce BCA Protein Assay Kit (ThermoFisher Scientific [23225]; Waltham, MA, USA) and final concentrations were adjusted to the same value for all samples prior to loading gels. For immunoblotting, equal amount protein samples were denaturated by incubation at 96^°^C for 8 m in loading buffer (pH 6.8) and loaded to SDS polyacrylamide gel for electrophoresis (SDS-PAGE) at 60V for 15 m and 100V for 2-3 h. Proteins in gels were then transferred to nitrocellulose blotting membrane (Cytiva [10600008]; Marlborough, MA, USA) at 19V for 65 m by a semi-dry transfer cell (Bio-Rad [1703940]; Hercules, CA, USA), following with a blocking step by incubation with a 5% skim milk – 1% bovine serum albumin (BSA) (ThermoFisher Scientific [BP1600-100]; Waltham, MA, USA) cocktail in 1X tris-buffered saline (TBS) with 0.1% (v/v) Tween 20 (TBST) for 1 hour. After the blocking step, all membranes were incubated overnight at 4°C with the specific primary antibody. The next morning, membranes were washed 3 times with TBST and then incubated at room temperature for 1 hour with anti-rabbit, anti-mouse, or anti-goat secondary antibodies conjugated to horseradish peroxidase (HRP) (Agilent Dako; Santa Clara, CA, USA) diluted at 1:5,000 in TBST. The membranes were washed 3 times with TBST again and incubated with enhanced chemiluminescent substrates for HRP (low picogram to mid-femtogram levels) (ThermoFisher Scientific; Waltham, MA, USA) before taking images for protein detection by using ChemiDoc Imager (Bio-Rad; Hercules, CA, USA). Proteins targeted selectively by a specific antibody were detected on the membrane according to their molecular weight estimated by a protein ladder (Bio-Rad [1610374]; Hercules, CA, USA). All images were captured with colorimetric and chemiluminescence channels, and these image types were merged for a more precise data production. Corresponding protein bands on the membranes were quantified from raw images by ImageJ (National Institutes of Health (NIH); Bethesda, MD, USA). The following primary antibodies were used in this study: Acyl-CoA thioesterase 9 (Acot9) [HPA035533], β-actin [A5441] (MilliporeSigma; St. Louis, MO, USA); ACOT9 [sc-100476], cytochrome C [sc-13560], heat shock protein 90 (HSP90) [sc-13119] (Santa Cruz Biotechnology; Dallas, TX, USA); NADH:ubiquinone oxidoreductase core subunit S7 (NDUFS7) (Bio-Techne (NOVUS) [NBP1-49846]; Minneapolis, MN, USA); HSP60 [12165S], pyruvate kinase 2 (PKM2) [4053T], PKM1/2 [3190T], pyruvate dehydrogenase (PD) [3205T], hexokinase I (HK I) [2024T], hexokinase II (HK II) [2867T], glyceraldehyde-3-phosphate dehydrogenase (GAPDH) [5174T] (Cell Signaling Technology; Danvers, MA, USA); Uncoupling protein 1 (UCP1) (Abcam [ab23841]; Cambridge, UK).

### Isolation and Cultures of Brown Preadipocytes

Precursor cells for primary brown adipocyte culture were isolated from 4-to-6-week-old *Acot9^B-KO^* and *Acot9^B-WT^*littermates and cultured as previously described^17^. In brief, mouse BATs were dissected, rinsed, and kept in cold PBS until all BAT pads were harvested on the day of the experiment. PBS was then replaced with the digestion media containing collagenase B (MilliporeSigma [SCR103]; St. Louis, MO, USA) and dispase II (MilliporeSigma [4942078001]; St. Louis, MO, USA), and BAT pads were minced before transferring the mix into a 50 ml tube (10 ml total volume of digestion media). Samples were digested by gently shaking on an orbital shaker for 25 m at 37°C. Digestion was stopped by adding 10 ml growth media DMEM/F12 (ThermoFisher Scientific [11320033]; Waltham, MA, USA) supplemented with 1% penicillin/streptomycin and 20% fetal bovine serum (FBS). Cells were filtered through 100 and 40 µm cell strainers into fresh 50 ml tubes and centrifuged at 300 x g for 5 m at room temperature before seeding for each study. Cells were then seeded on collagen-coated plates (Bio-Techne [3443-100-01]; Minneapolis, MN, USA). The growth media were changed every day until cells became confluent. Upon reaching confluency, cells were induced for adipogenic differentiation with the differentiation media (DMEM/F12 with 1% penicillin/streptomycin, 10% FBS, 5 µg/ml insulin (MilliporeSigma [I0516]; St. Louis, MO, USA), 0.5 mM 3-isobutyl-1-methylxanthine (IBMX) (MilliporeSigma [I5879]; St. Louis, MO, USA), 1 µM rosiglitazone (MilliporeSigma [R2408]; St. Louis, MO, USA), 5 µM dexamethasone (MilliporeSigma [D4902]; St. Louis, MO, USA), 0.125 mM indomethacin (MilliporeSigma [I7378]; St. Louis, MO, USA), 1 nM T3 (MilliporeSigma [T5516]; St. Louis, MO, USA), 2 µM tamoxifen (MilliporeSigma [T5648]; St. Louis, MO, USA)). Cells were replenished with the differentiation media for the first 2 days and switched to maintenance media (DMEM/F12 with 1% penicillin/streptomycin, 10% FBS, 5 µg/ml insulin hormone (MilliporeSigma [I0516]; St. Louis, MO, USA), 1 µM rosiglitazone (MilliporeSigma [R2408]; St. Louis, MO, USA), 1 nM T3 hormone (MilliporeSigma [T5516]; St. Louis, MO, USA), 2 µM tamoxifen (MilliporeSigma [T5648]; St. Louis, MO, USA)) for the next 6 days (until end of the 8^th^ day).

### Cultured Primary Brown Adipocyte Metabolic Analyses

Oxygen consumption rates (OCR) and ECAR were determined in *Acot9^−/−^* and WT primary brown adipocytes by using a Seahorse XFe96 Analyzer (Agilent; Santa Clara, CA, USA). Isolated precursor cells were seeded at 2,000/well into precoated XF96 Seahorse plates and induced differentiation. From the 6^th^-to-8^th^ days of differentiation, primary brown adipocytes were treated with 1 µM L-(-)-NE (+) bitartrate (MilliporeSigma [Calbiochem 489350]; St. Louis, MO, USA) supplemented in the maintenance media. On day 7, Seahorse cartridges were rehydrated with the calibrant solution overnight in a non-CO_2_ incubator at 37°C. Prior to the assay on day 8, injection ports on the sensor cartridge were loaded as follows: Port A – 20 µl of NE (final concentration 5 µM in the well), port B 22 µl of NucRed Live 647 (ThermoFisher Scientific [R37106]; Waltham, MA, USA). Plated cells were rinsed twice with 200 µl assay media (Krebs-Henseleit buffer (pH 7.4) with 0.45 g/L glucose, 111 mM NaCl, 4.7 mM potassium chloride (KCl), 2 mM MgSO4^-^7H_2_O, 1.2 mM Na2HPO4, 5 mM HEPES, and 0.5 mM carnitine), added assay media for a final 180 µl/well volume, and incubated in a non-CO_2_ incubator at 37°C for 1 hour before the assay. OCR and ECAR were measured before and after the port A injection for NE (1 µM), and data were normalized with total live adipocyte count per well.

### Oil-red o Staining

To measure neutral (total) lipid accumulation in *Acot9^−/−^* and *WT* primary brown adipocytes, cells were washed 3 times with PBS, and then fixed by adding 10% formalin for 1 h at 22°C. Oil-red O (ORO) working solution was prepared by mixing 30 ml of ORO stock solution (MilliporeSigma [O1391]; St. Louis, MO, USA) with 20 ml water and filtered through a filter funnel. Fixed cells were first incubated in 600 µl 60% isopropanol and then 2 ml ORO working solution for 5 m for each step. Cells were then rinsed with tap water until water ran clear and kept wet until the histology/densitometry analysis. For optical densitometry (O.D.) measurement, cells were incubated with 600 µl isopropyl alcohol for 15 m and collected isopropyl alcohol extract were measured at 492 nm using a SpectraMax i3x spectrometer (Molecular Devices; San Jose, CA, USA).

### Fatty acid uptake

To measure FA uptake in *Acot9^−/−^* and *WT* primary brown adipocytes, radiolabeled palmitic acid was used as substrate as previously described^40,74^. For incubation, 1 μCi [^14^C]-palmitate (American Radiolabeled Chemicals Inc.; St. Louis, MO, specific activity: 55 μCi/μmol) supplemented with 200 μM cold palmitate was conjugated in 5:1 molar ratio with FA free BSA (MilliporeSigma [A8806]; St. Louis, MO, USA). Cultured cells were first incubated in a serum-free maintenance media for 2 h, followed by an incubation with maintenance media containing labeled FA for a short period (30 s), and the media were immediately aspirated after that. The culture plate was placed on ice, and cells were washed 3 times with ice-cold PBS. Cells were then scraped from the surface with 200 µl PBS and homogenized with a pestle motor. Radioactivity in homogenates were measured by transferring 150 µl samples into 4 ml Ecoscint H using a LS6000IC liquid scintillation counter (Beckman Coulter Inc.; Brea, CA).

### Fatty acid oxidation

FA oxidation (FAO) rate was measured by the oxidation of [^14^C]-palmitate into [^14^C]-labeled CO_2_ ([^14^C]-CO_2_) and [^14^C]-labeled acid soluble metabolites ([^14^C]-ASM) in primary brown adipocytes that were differentiated in 60 mm dishes and cultured in maintenance media for seven days prior to the FAO assay. Cells were washed 3 x with warm PBS (37°C) and incubated in serum-starvation media (M199 media supplemented with 1% penicillin/streptomycin) for 2 h. Cells were then incubated with 1 ml serum-starvation media supplemented with 1 mM carnitine and 200 µM [^14^C]-palmitate complexed to BSA (5:1 molar ratio; 1 μCi/ml) for 3 h at 37°C. 250 µl culture media was mixed with 200 µl of 1 M perchloric acid in a 1.5 ml tube. A filter paper disc was fixed to the cap and humidified with 20 µl of 1 M NaOH. The samples were shaken for 1 hour at room temperature without contact with the filter paper. Complete palmitate oxidation into CO_2_ was captured by [^14^C]-labeled CO_2_ that was trapped in the filter paper. The filter paper was placed in 4 ml Ecoscint H and activity measured by scintillation counter as mentioned previously. Incomplete oxidation was detected in ASM by separating the insoluble aggregate by centrifugation (14,000 x g for 10 m at 4°C) and measuring the activity within the soluble supernatant after mixing it into 4 ml Ecoscint H. Radioactivity was normalized to the specific activity of [^14^C]-palmitate (55 μCi/μmol) and protein concentration within each dish.

### *De novo* lipogenesis

Rates of *de novo* lipogenesis in *Acot9^−/−^* and WT primary brown adipocytes were measured as follows: fully-differentiated cells were serum-starved for 2 h in 1 ml M199 media supplemented with 1% penicillin/streptomycin and washed 3 x 1 ml warm PBS. Cells were then incubated with serum-free and glucose-free culture media (DMEM) that contain 1 mM (1 μCi/ml) [^14^C]-acetate (PerkinElmer; Waltham, MA, USA; specific activity: 50 μCi/μmol) or 13.5 mM (1 μCi/ml) [^14^C]-glucose PerkinElmer, specific activity: 250-360 μCi/μmol) at 37°C for 3 and 4 h, respectively. At the end of the incubation period, cells were moved onto ice and washed 3 x 1 ml cold PBS. Cells were eventually scraped into 300 µl cold PBS and homogenized in 1.5 ml tubes using a plastic (motor-driven) pestle. A 50 µl sample was aliquoted for each homogenate for protein quantification. A 200 µl sample was then transferred into a conic lipid extraction tube containing 0.5 ml methanol and 1 ml chloroform. Solutions were vortexed for 2 m, and centrifuged (3,000 x g for 15 m at 22°C). The organic phase for each sample were transferred into a new tube, added 1 ml chloroform to the remaining solution, and centrifuged (3,000 x g for 15 m at 22°C). The organic phases were combined and dried under nitrogen gas flow. The lipids in the dry pellets were resuspended in 300 µl chloroform, and after a quick vortex, 100 µl of the samples were transferred into a vial containing 4 ml Ecoscint H for measurements using a LS6000IC liquid scintillation counter (Beckman Coulter Inc.; Brea, CA).

### *In vitro* lipolysis

Lipolytic activity was determined as a function of glycerol release from glycerolipids. Briefly, mouse primary brown adipocytes were serum-starved overnight (16 h) in DMEM without FBS. Cells were then incubated in lipolysis assay buffer comprising DMEM without glucose/glutamine/phenol red (ThermoFisher Scientific [A1443001]; Waltham, MA, USA) supplemented with 1 mg/ml glucose, 2% fatty acid-free BSA with or without 1 nM isoproterenol for 3 h. Glycerol within the incubation media was measured using an MAK215 lipolysis colorimetric assay kit (MilliporeSigma; St. Louis, MO, USA). Glycerol release was normalized to the total protein content.

### Glucose uptake

*Acot9^−/−^* and WT primary brown adipocyte glucose uptake rates were determined using a Glucose-Glo Assay (Promega [J6021]; Madison, WI, USA) according to the manufacturer’s instructions. In short, cellular glucose uptake was measured using a stable glucose analog, 2-deoxyglucose (2-DG). Cells that were fully differentiated in a 96-well plate were washed with PBS to remove glucose from the culture media. 50 µl of freshly prepared 1 mM 2-DG were added into each well and incubated for 10 m at 22°C. The uptake process was halted by adding 25 µl Stop Buffer with a gentle shake. Cells were then supplemented with 25 µl of Neutralization Buffer followed by 100 µl of 2-DG6P Detection Reagent. Solutions were then incubated at 22°C for at least 30 m and analyzed by luminescence using a SpectraMax i3x spectrometer (Molecular Devices; San Jose, CA, USA) for 0.3-to-1 second integration.

### *Ex vivo* lipolysis

HFD-fed WT and *Acot9^−/−^* mice were sacrificed after a 6-hour fast. Gonadal fat pads were dissected and cut into 50 mg pieces for incubation at 37°C in 1 ml of phenol red-free DMEM containing 2% fatty acid-free BSA with or without 1 μM isoproterenol for 2 h. Accumulation of non-esterified fatty acids (NEFA) and glycerol within the incubation media was measured using MAK211 lipolysis colorimetric assay kit (MilliporeSigma; St. Louis, MO, USA) for glycerol and a diagnostic assay kit (FUJIFILM Wako Diagnostics; Mountain View, CA, USA) for NEFA. Lipolytic releases were normalized to the tissue weight.

### RT-qPCR

Total RNA was extracted by TRIzol-based methodology according to the manufacturer’s instructions, and corresponding cDNA was synthesized using the SuperScript III Reverse Transcriptase (ThermoFisher Scientific [18080044]; Waltham, MA, USA). Transcript abundance was measured using SYBR Green PCR Master Mix (ThermoFisher Scientific [4309155]; Waltham, MA, USA) in a LightCycler 480 System (Roche Diagnostics; Indianapolis, IN, USA). Relative Ct values were normalized to the geometric mean of 3 housekeeping genes: GAPDH (glyceraldehyde 3-phosphate dehydrogenase), TBP (TATA box binding protein), and RPL32 (ribosomal protein L32) using the 2^-▴▴Ct^ method. Primer sequences are listed Supplementary Table 1.

### Recombinant ACOT9 protein

Plasmids (pMAL-c2X [#75286]) containing the sequence for the maltose binding protein (MBP)-ACOT9 fusion protein were generated and kindly provided by Drs. Veronika Tillander and Stefan E. H. Alexson (Karolinska Institutet, Sweden)^40^. For MBP containing negative control, Acot9 sequence was removed from the plasmid by XbaI restriction (New England BioLabs [R0145S]; Ipswich, MA, USA) followed by the quick ligation of the open plasmid by the Quick Ligation Kit (New England BioLabs [M2200]; Ipswich, MA, USA). For expression of MBP-Acot9, 50 µl of BL21 (DE3) bacteria (New England BioLabs [C2527H]; Ipswich, MA, USA) was pipetted into a 1.5 ml tube, and added 50 ng equivalent of plasmids. The cell mixture was carefully flicked and incubated on ice for 30 m, following a quick heat shock for 10 s at 42°C, and immediately placed on ice again. The mixture was then diluted with SOC outgrowth media (New England BioLabs [B9020S]; Ipswich, MA, USA) and incubated by shaking vigorously at 37°C for 60 m. Bacteria solution was spread on selection plates with Luria-Bertani (LB) media (+100 µl/ml ampicillin) and incubated overnight at 37°C. Untransformed BL21 (DE3) bacteria was used as a negative control. In 14 ml round-bottom polypropylene tubes, 4 ml LB media (liquid) were inoculated with the BL21 colony from the selection plate and rotated at 37°C for 4 h. 1 ml of cell mixture was used to inoculate 100 ml of LB media without antibiotics and rotated at 37°C. At 600 nm, O.D. was measured every 2 h to determine optimal bacterial growth condition for the induction of protein production. At the O.D. of 0.5, protein production was initiated by adding 1M isopropyl β-D-1-thiogalactopyranoside (IPTG) and rotating overnight at room temperature. Bacteria were then transferred into 2 x 50 ml tubes and centrifuged at 4,000 x g for 3 m at 4°C. Following the removal of LB media, the pellets were homogenized in 5 ml cold BugBuster Protein Extraction Reagent (MilliporeSigma [70584]; St. Louis, MO, USA), and sonicated (30% 15 s twice/sample) to break open the bacteria. Homogenates were centrifuged at 14,000 x g for 15 m at 4°C, after which the supernatant was further centrifuged at 100,000 x g for 30 m at 4°C. The supernatant containing the recombinant MBP-ACOT9 or MBP control were aliquoted.

### Maltose-binding protein pulldown assay with amylose beads

Equal amount of crude MBP-ACOT9 and MBP proteins were thawed and incubated with 2 mg amylose magnetic beads (AMB) (New England BioLabs [E8035S]; Ipswich, MA, USA) at 4°C in rotation for 1 h. The AMB-bound proteins were precipitated with a magnetic rack, washed 3 times with cold PBS, and resuspended in 1 ml RIPA buffer with protease and phosphatase inhibitor cocktails. The pulldown experiments were conducted using protein lysate (1 mg/ml) that were collected from human embryonic kidney (HEK) 293E cells. 1 ml of HEK293E protein lysate was incubated with 100 µl of MBP-ACOT9 and MBP proteins bound to magnetic beads. Because protein-protein interactions of hot-dog fold domains varies with temperature, the incubation was carried out by rotating first at 4°C overnight followed by 1 hour at 22°C and 30 m at 37°C. Following the incubation, AMBs and co-pulled proteins were washed 3 x 5 m with cold PBS using a magnetic rack. Proteins were eluted with 25 µl loading buffer and analyzed in immunoblots as described above.

### *In situ* proximity ligation assay

Proximity ligation assay was carried out according to the manufacturer’s instructions (MilliporeSigma [DUO92101]; St. Louis, MO, USA). Initially, cover glasses were coated in 24-well plates containing 5 µg/ml polyethyleneimine for 1 hour at 22°C. HEK293E cells were seeded at 40% confluency, and the next day, they were treated for 15 m with either 1 μM FCCP or dimethyl sulfoxide (v/v) as control. Cells were then washed with ice-cold PBS and fixed in 4% paraformaldehyde (PFA) for 10 m at 22°C. To prevent any background staining, cells were incubated with Duolink Blocking Solution (40 µl/cover glass) in a heated humidity chamber (1 hour at 37°C). Cells were then incubated overnight at 4°C with the primary antibody solution (40 µl/cover glass), containing antibodies against ACOT9 (Santa Cruz Biotechnology [sc-100476]; Dallas, TX, USA) and NDUFS7 (Proteintech [15728-1-AP]; Rosemont, IL, USA), diluted 1:100 in Duolink Antibody Diluent. Slides were washed (3 x 5 m) with Duolink Wash Buffer followed by incubation (1 hour in a preheated humidity chamber at 37°C) with the secondary antibody solution containing 8 µl of PLA probe MINUS stock, 8 µl of PLA probe PLUS stock, and 24 µl of Antibody Diluent (40 µl/cover glass). Slides were then washed with Duolink Wash Buffer A (3 x 5 m at 22°C) and incubated with Duolink Ligase in a preheated humidity chamber (30 m at 37°C). Following the ligation, the steps of amplification, mounting and imaging were completed in the dark. For the amplification, slides were first washed in Duolink Wash Buffer A (3 x 5 m at 22°C) followed by incubation in a preheated humidity chamber (30 m at 37°C) with the amplification mix containing Polymerase diluted 1:80 in Duolink Amplification Buffer (40 µl/cover glass). For the final washes, slides were first washed (2 x 10 m at 22°C) in 1X Wash Buffer B followed by a wash in 0.01X Wash Buffer B (10 m at 22°C). Cover slides were then mounted with 8 µl SlowFade Diamond Antifade Mounting Medium with DAPI (4,6-diamidino-2-phenylindole) (ThermoFisher Scientific [S36973]; Waltham, MA, USA). Images were taken by Axioscan 7 automated scanning microscopy (Carl Zeiss Inc.; White Plains, NY, USA) for DAPI and Texas Red.

### Analysis of thioesterase activity

BAT whole tissue lysates from HFD-fed *Acot9^B-KO^* and *Acot9^B-WT^* mice were used to determine acyl-CoA thioesterase activity in mice. Accordingly, samples were diluted in freshly made prewarmed reaction buffer (pH 7.4) containing 50 mM KCl, 10 mM HEPES and 0.3 mM DTNB (Ellman’s reagent; MilliporeSigma [D8130]; St. Louis, MO, USA). The thioesterase activity was measured by detection of freed CoA-DTNB at 412 nm in every 20 s using a SpectraMax i3x spectrometer (Molecular Devices; San Jose, CA, USA). The commercially available acyl-CoA esters for exogenous substrate were purchased from MilliporeSigma (St. Louis, MO, USA). An E_412_=13,600 M^-1^ cm^-1^ (molecular extinction coefficient) was used for activity calculation following with enzymatic kinetics by nonlinear regression.

### Statistical analyses, figure generation, and reproducibility

Experiments were conducted after mice were randomized to the groups. All measurements and analyses were confirmed more than once. Group sizes were determined by power analyses prior to each experiment calculating the sample size at 80% beta and 0.05% alpha and expected values for each assay. Cell culture experiments were repeated in at least three independent replicates. Investigators were not blinded to experimental groups. Image analyses and densitometry were done via blinded evaluation for unbiased analyses. The clinical data was tested and confirmed by a biostatistician (XKZ). For diet-matched genotype comparisons, independent samples t-test was used. Thermography data was analyzed by detecting changes in the slope of a linear regression fit. Group comparisons with multiple variables were analyzed for significance by repeated measures two-way analysis of variance (ANOVA), followed by Bonferroni’s post hoc analysis. Statistical analyses and plotting of data was done using GraphPad Prism (Dotmatics; Bishop’s Stortford, UK), Excel Software (Microsoft Corporation; Redmond, WA, USA), VassarStats for ANCOVA analysis, MetaboAnalyst 5.0 online tool for pathway analysis of the metabolomics data, and R programming language by R Core. Illustrations were generated with BioRender.

## Acknowledgments

We thank Geoffrey C. Farrell (The Australian National University, Australia), Isabelle A. Leclercq (Université Catholique de Louvain, Belgium), and Paul Cohen (The Rockefeller University, USA) for insightful comments on the manuscript. We also thank David E. Cohen (Brigham and Women’s Hospital, Harvard University, USA) and his laboratory members for helpful discussions. We thank Timothy E. McGraw (WCM) for providing the *Ucp1-Cre* mouse, and Veronika Tillander and Stefan E. H. Alexson (Karolinska Institutet, Sweden) for providing the recombinant ACOT9 plasmid construct. We thank Frederick R. Maxfield and Timothy A. Ryan (WCM) for their expert advice on the biochemistry of protein-protein interaction. Finally, we thank Weill Cornell Medicine Metabolic Phenotyping Core, Proteomics and Metabolomics Core, and Research Animal Resource Center for excellent assistance with experiments.

## Author Contributions

FH expanded upon the concept, conceived the experimental plan, designed and performed experiments, analyzed data, and prepared the figures. JMJ analyzed proteomics and human transcriptomics data, conceived the interactome analysis, performed computational modeling and prepared the figures and tables. BDS designed and performed mass spectrometry-based proteomics and analyzed data. SS performed experiments. LL performed additional computational modeling of ACOT9-NDUFS7 complex interactions. VD contributed to thermography imaging. JQ produced recombinant protein. SSS performed proximity ligation assay. XKZ contributed to the analysis of the human RNAseq dataset. AJD provided the human RNAseq dataset. NMI collected human samples. HY oversaw ACOT9-NDUFS7 computational modeling. LCC oversaw mass spectrometry-based proteomics. BAE conceived the hypothesis, designed experiments, analyzed data, and directed the project. FH, JMJ, and BAE wrote the manuscript with input from all authors.

## Data and Materials Availability

All data that support the findings of this study are available in the manuscript or the supplementary materials, and available upon request. All scripts used in this study are available upon request.

## Supplementary Information is available for this paper

Correspondence and requests for materials should be addressed to Baran Ersoy. Inquiries specifically regarding the computational analyses should be directed to James M. Jordan.

Peer review information includes the names of reviewers who agree to be cited and is completed by Nature staff during proofing.

Reprints and permissions information is available at www.nature.com/reprints.

