## Extended Data Figures for "Submitochondrial Protein Translocation in Thermogenic Regulation"

[1]These authors contributed equally
[2]Division of Gastroenterology and Hepatology, Weill Center for Metabolic Health, Joan & Sanford I. Weill Department of Medicine, Weill Cornell Medicine; New York, NY 10021, USA
[3]Florida State University, Department of Biological Science; Tallahassee, FL 32304, USA
[4]Sandra and Edward Meyer Cancer Center, Joan & Sanford I. Weill Department of Medicine, Weill Cornell Medicine; New York, NY 10065, USA
[5]Division of Biostatistics, Department of Population Health Sciences, Weill Cornell Medicine; New York, NY 10065, USA
[6]Breast Medicine Service, Memorial Sloan Kettering Cancer Center; New York, NY 10065, USA
[7]Weill Cornell Medical Center; New York, NY 10065, USA
[8]Weill Institute for Cell and Molecular Biology and Department of Computational Biology, Cornell University; Ithaca, NY 14853, USA
[9]Center for Innovative Proteomics, Cornell University; Ithaca, NY 14853, USA
[10]Dana Farber Cancer Institute, Harvard Medical School; Boston, MA 02215, USA
[11]This author is now retired.

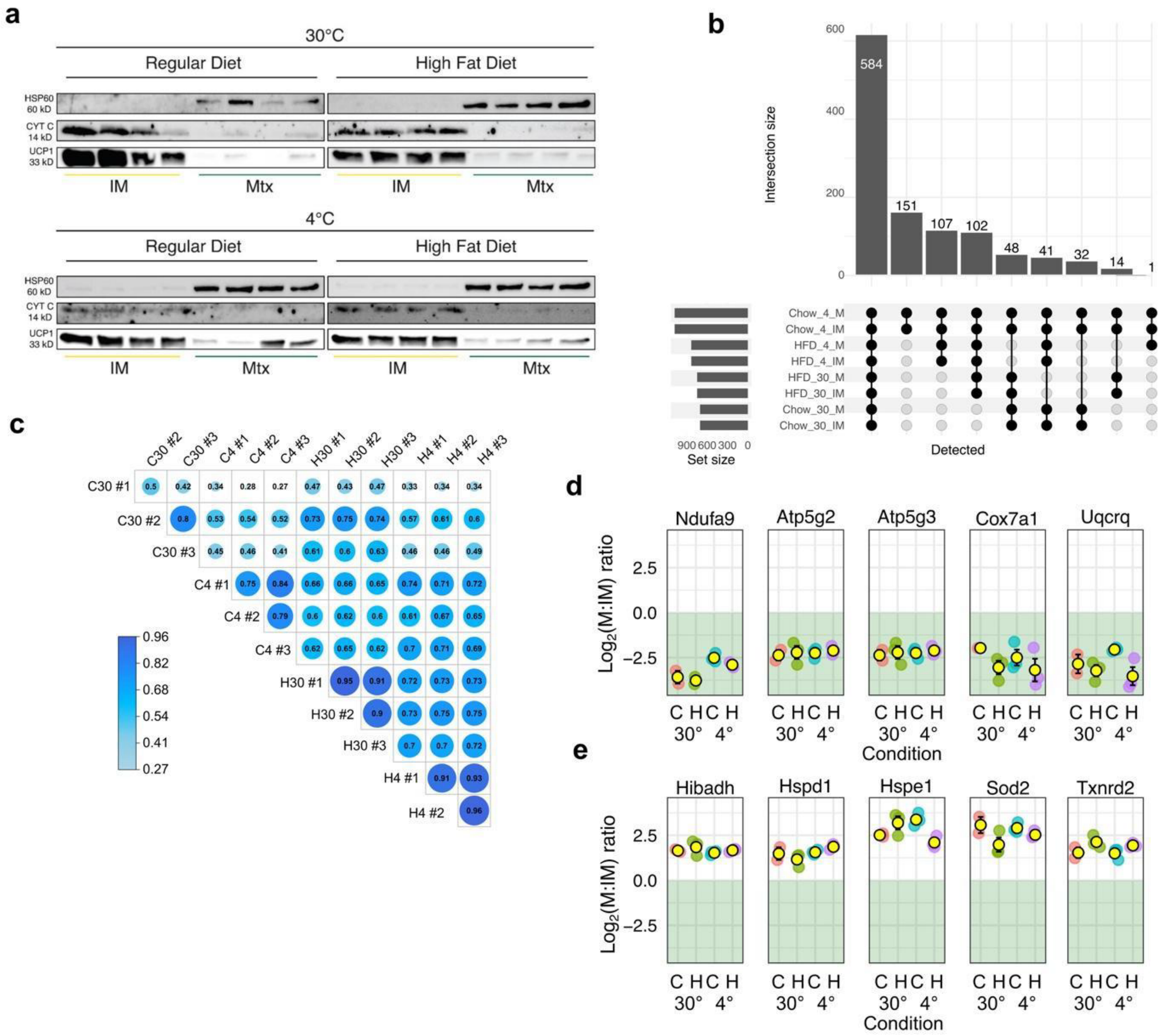

**Extended Data Fig. 1|Validation of submitochondrial proteomic approach and analysis.** **a**, Immunoblot of matrix and innermembrane (IM) markers with protein from cold and high-fat diet stressed brown adipose tissue (BAT) mitochondria. **b**, Upset plot of proteins detected in each fraction of each condition. **c**, Correlation of IM-matrix ratios within and without experimental conditions. **d**, Ratios of known IM-anchored and **e**, matrix-localized proteins. For d and e, each dot is from an individual mouse sample. Yellow dot with an outline is the mean. Error bars indicate ± SEM. Green shading indicates IM enrichment, and an absence of shading indicates matrix enrichment.

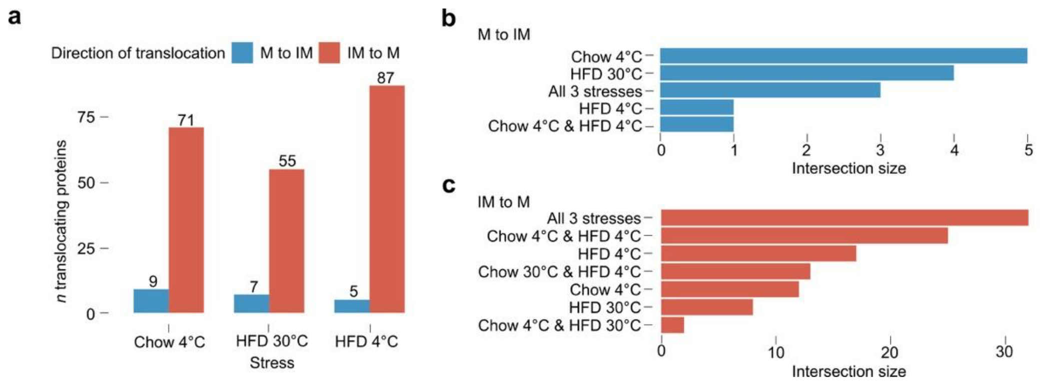


**Extended Data Fig. 2|Identification of perturbation-specific and perturbation-nonspecific submitochondrial translocators. a**, Number of genes which fit the criteria for switching submitochondrial localization in different stress conditions. **b,** Overlapping set analysis of proteins which meet criteria for IM-to-matrix switching or **c**, matrix-to-IM switching in cold, high fat diet (HFD), or a combination of both cold and HFD compared to control animals housed at thermoneutrality and fed a standard chow diet.

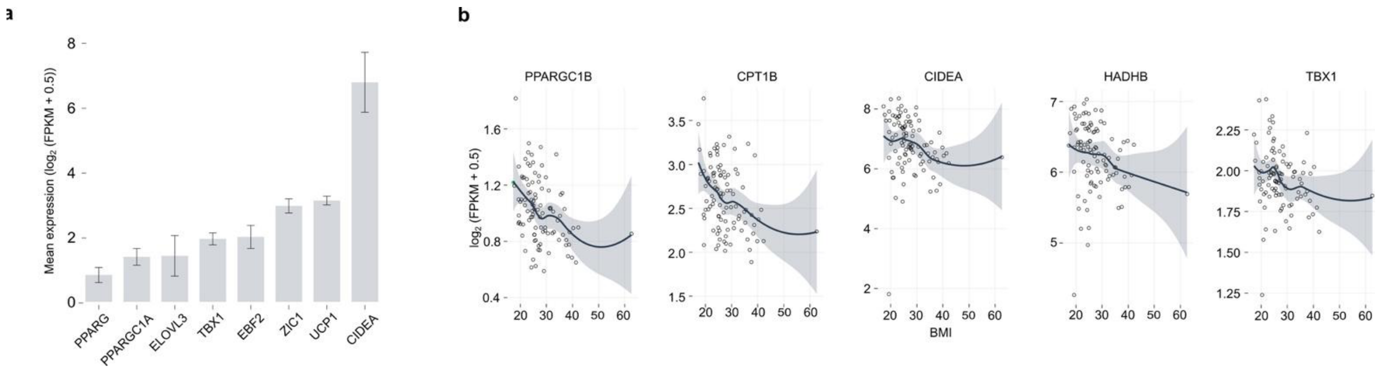


**Extended Data Fig. 3|Thermogenic gene expression levels in scWAT of 100 patients of varying BMI. a**, Mean expression levels of thermogenic genes in 100 human scWAT transcriptomes. Genes with mean $\log_2$ (FPKM+0.5) > 0 considered expressed. Error bars indicate +/- SEM. **b,** Scatter plots showing the relationship between BMI and gene expression for each thermogenic gene with significant negative correlation. FDR < 0.01, ***FDR < 0.01, rho > 0, Spearman's test The line represents the LOESS smoothed fit with 95% confidence interval ribbons.

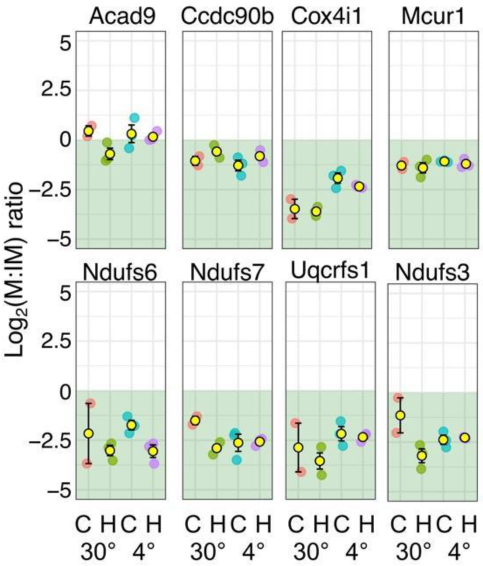


**Extended Data Fig. 4|Enrichment of IM-anchored subunits with high degree connectivity.** Each dot is from an individual mouse sample. Yellow dots indicate the mean. Error bars indicate ± SEM. Green shading indicates IM enrichment, and an absence of shading indicates matrix enrichment.

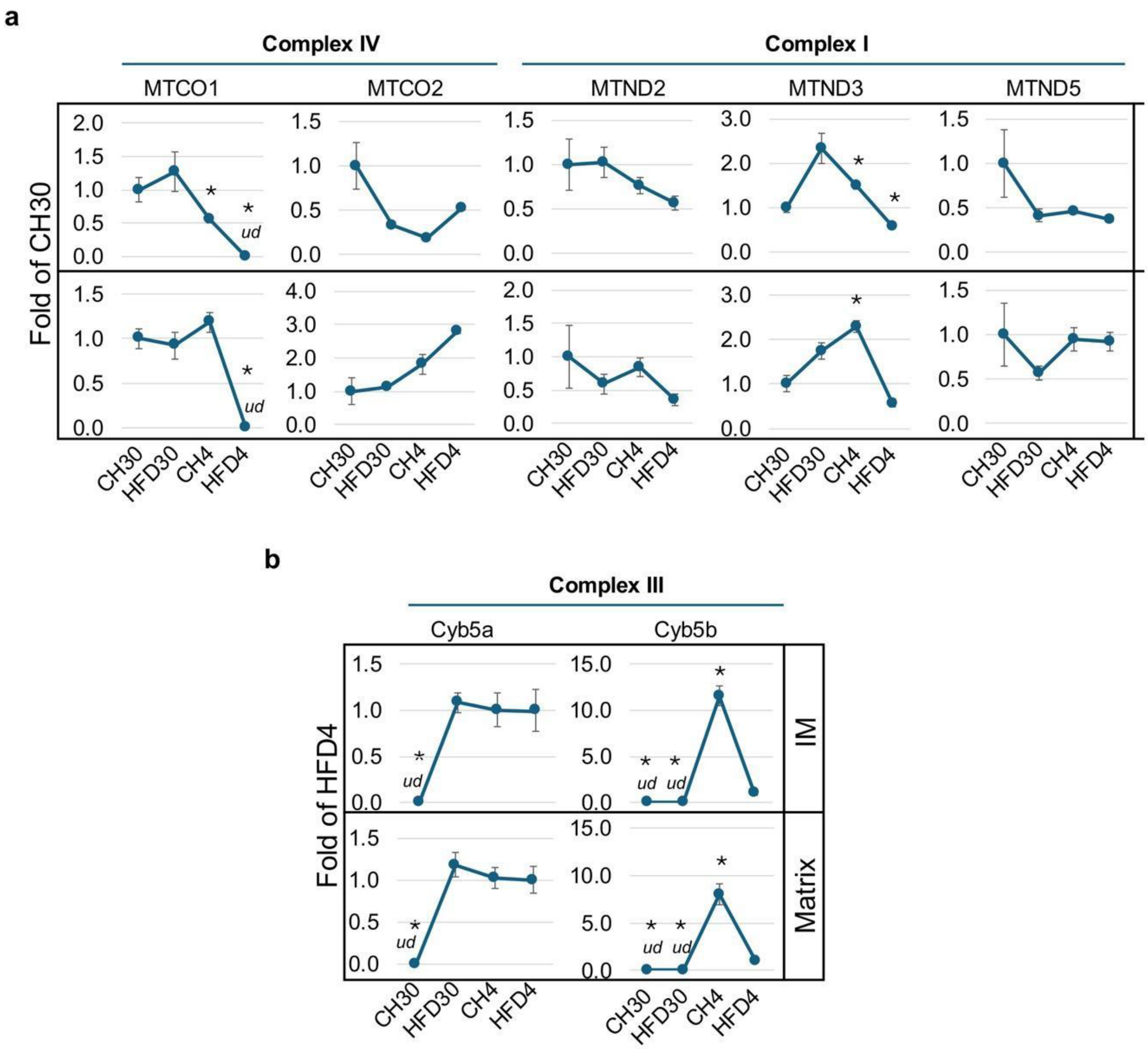


**Extended Data Fig. 5|Complex effects on mitochondrially-encoded mitoribosome-dependent gene expression. a**, Inconsistent effect on mitoribosome-target gene expression levels seen in ETC I and IV subunits and **b**, ETC III-associated electron carriers. For a and b, n=3 animals/condition; *P-value < 0.05, independent samples t-test. *ud*: undetected.

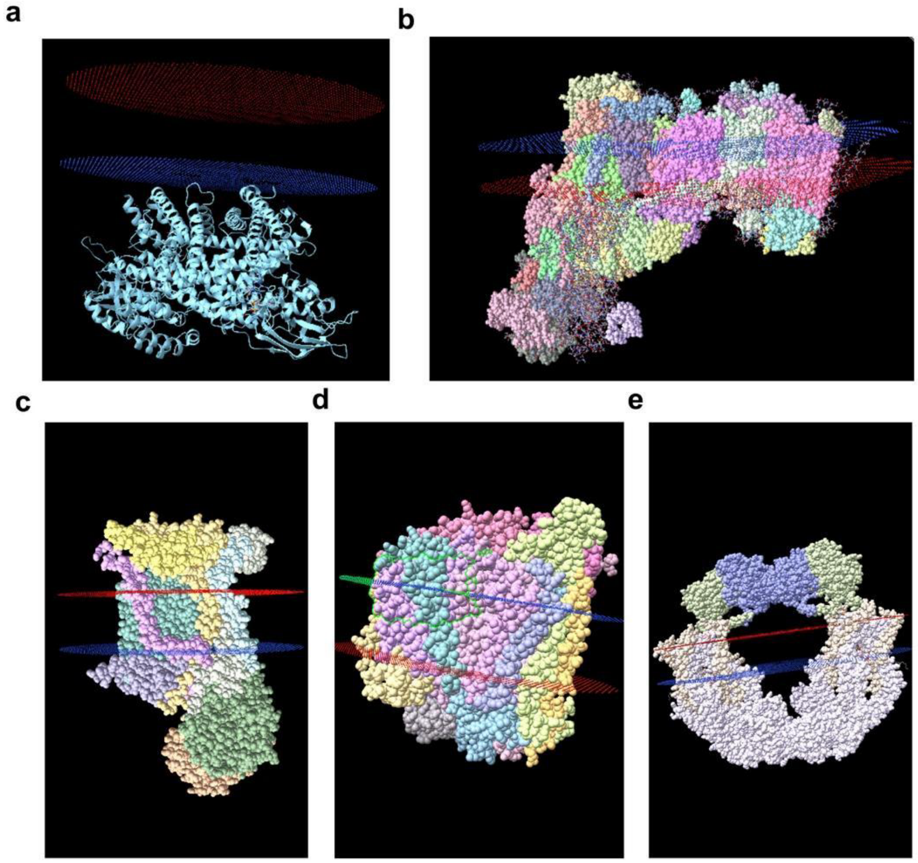


**Extended Data Fig. 6|PPM 3.0 models of solved IM complexes embedded in a simulated mitochondrial IM phospholipid bilayer to identify matrix-exposed histidine residues. a**, ACAD9 homodimer (8PHF), **b**, ETC I (5LDW), **c**, ETC III (5XTE), **d**, ETC IV (5Z62), and **e**, MCUC (6K7Y) computationally embedded in IM using PPM3.0 and visualized with ChimeraX.

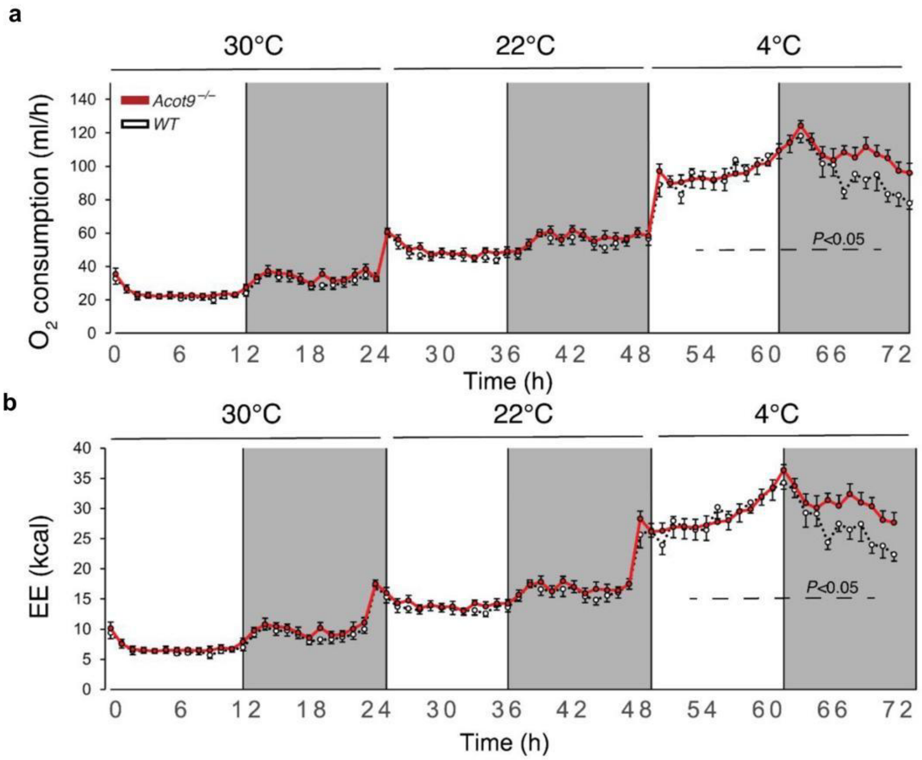


**Extended Data Fig. 7|Oxygen consumption and energy expenditure is increased in HFD-fed *Acot9*$^{-/-}$ mice.** **a**, Indirect calorimetric analysis of $VO_2$ and **b**, EE in HFD-fed *Acot9*$^{-/-}$ and WT mice (n=6) at thermoneutrality (30°C), room temperature (22°C), and cold exposure (4°C). Mice were acclimated to each temperature drop condition for two weeks prior to the measurements. Data represent hourly mean ± SEM; by two-way analysis of variance (ANOVA) with Bonferroni's post hoc correction (adjusted $P<0.05$).

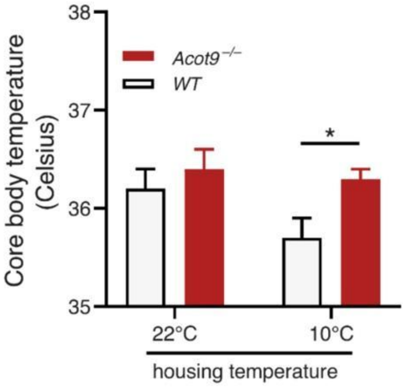


**Extended Data Fig. 8|Core body temperatures of *Acot9*$^{-/-}$ and WT mice at different housing temperatures.** HFD-fed Acot9$^{-/-}$ and WT mouse body temperatures (n=6/genotype) measured by a rectal thermometer at room temperature and two weeks following mild cold exposure (10°C). Graphs show mean + SEM; analyzed by one-way analysis of variance (ANOVA) with Bonferroni's post hoc correction (*P<0.05).

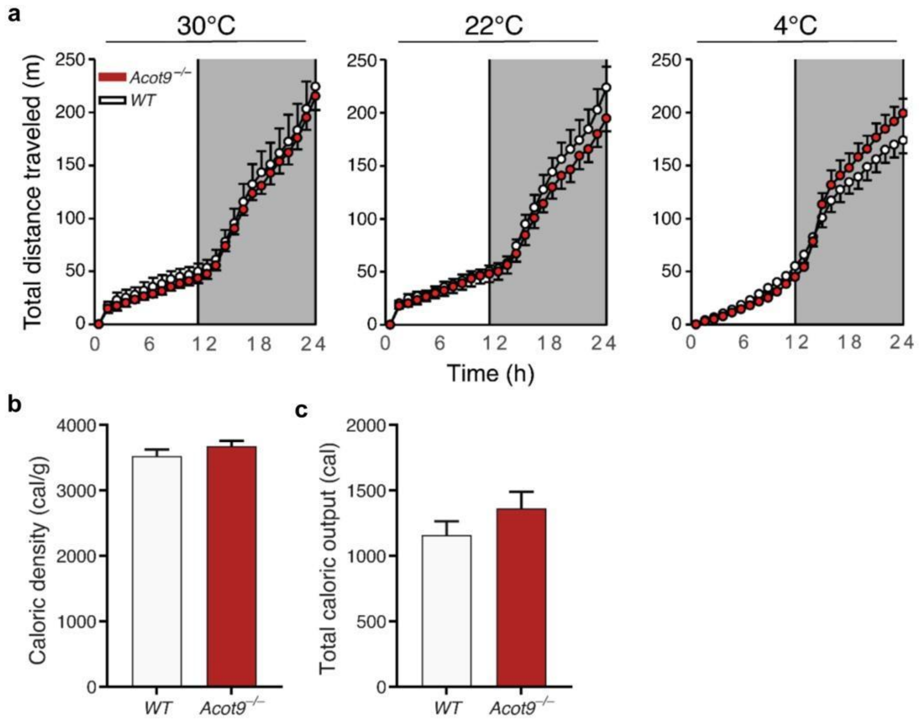


**Extended Data Fig. 9|Locomotor activity and nutrient absorption in HFD-fed *Acot9*$^{-/-}$ and WT mice.** **a**, Cumulative travel distances of HFD-fed *Acot9*$^{-/-}$ and WT mice (n=6/genotype) measured at thermoneutrality, room temperature, and cold exposure during indirect calorimetry measurements. **b**, Caloric density and **c**, gross calorific value of fecal samples. n=8-10/genotype. Plots show mean ± SEM.

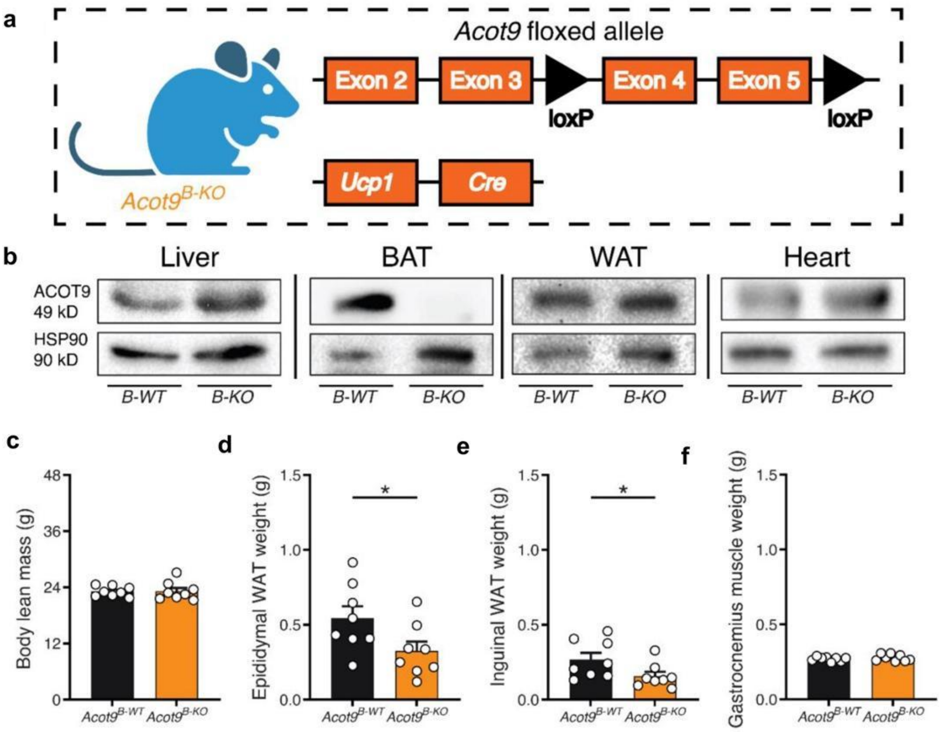


**Extended Data Fig. 10|Generation of BAT-specific Acot9 knock-out model and initial phenotyping. a**, Schematic of the tissue-specific transgenic knock-in of loxP sites and BAT-specific UCP1-driven *cre* recombinase. **b**, ACOT9 (and HSP90 control) immunoblot from multiple tissues sampled from HFD-fed mice. **c**, EchoMRI measurement of total lean mass in HFD-fed mice. **d**, Epididymal, **e**, inguinal white adipose tissues (WATs), and **f**, gastrocnemius muscle weights in HFD-fed mice. For c-f, n=8/genotype, circles indicate a measurement from an individual animal. Bars indicate mean. Error bars indicate + SEM; analyzed by independent samples t-test (*P<0.05).

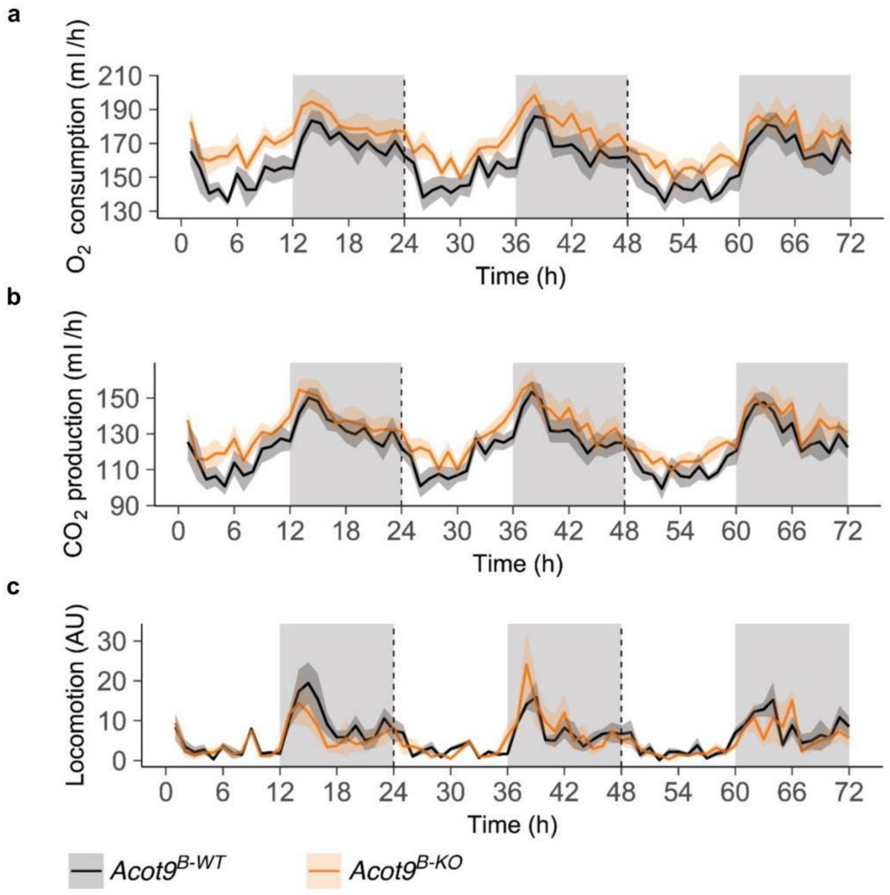


**Extended Data Fig. 11|Indirect calorimetry analysis of HFD-fed *Acot9*$^{B\text{-}KO}$ and *Acot9*$^{B\text{-}WT}$ mice.** **a**, Oxygen consumption ($VO_2$), **b**, $CO_2$ production ($VCO_2$), and **c**, physical activity levels over 3 d period in HFD-fed *Acot9*$^{B\text{-}KO}$ and *Acot9*$^{B\text{-}WT}$ mice. For a-c, n=5 mice, line indicates the average across all animals and ribbons indicate 95% confidence intervals.

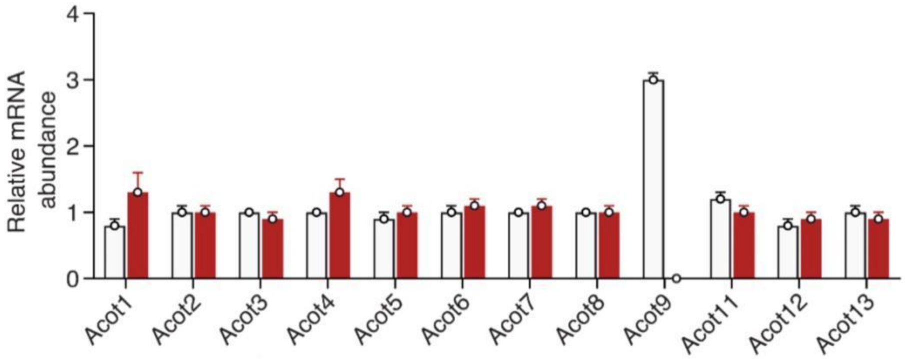


**Extended Data Fig. 12|Lipid metabolism gene expression in HFD-fed *Acot9*$^{-/-}$ BAT.** RT-qPCR analyses of ACOTs in BAT of HFD-fed *Acot9*$^{-/-}$ and WT mice (n=5/genotype). Relative abundances were determined by normalizing to the mean of three housekeeping genes: Gapdh (glyceraldehyde 3-phosphate dehydrogenase), Tbp (TATA box binding protein), and Rpl32 (ribosomal protein L32). Bars indicate mean and error bars indicate + SEM. *P < 0.05, **P < 0.01, independent samples t-test.

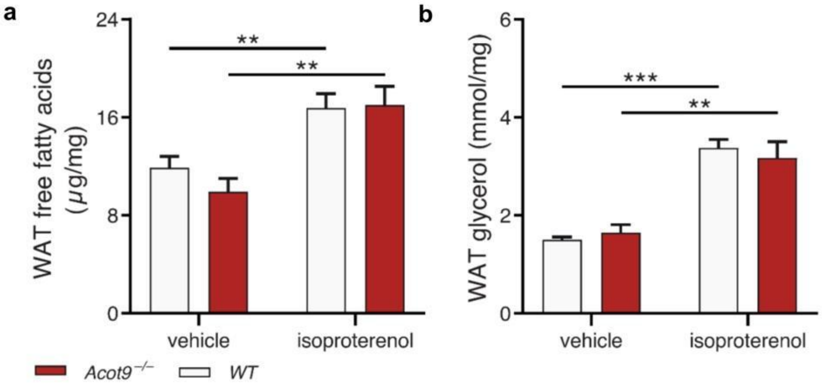


**Extended Data Fig. 13|Acot9 expression does not affect *ex situ* lipolysis in WAT. a**, Lipolysis rates were measured in WAT of HFD-fed Acot9$^{-/-}$ and WT mice after 6 h fast by quantifying non-esterified fatty acids and **b**, glycerol accumulation with 1 μM isoproterenol or vehicle (DMSO). For a and b, bars indicate the mean and error bars indicate + SEM; ns = not significant, analyzed by independent samples t-test.

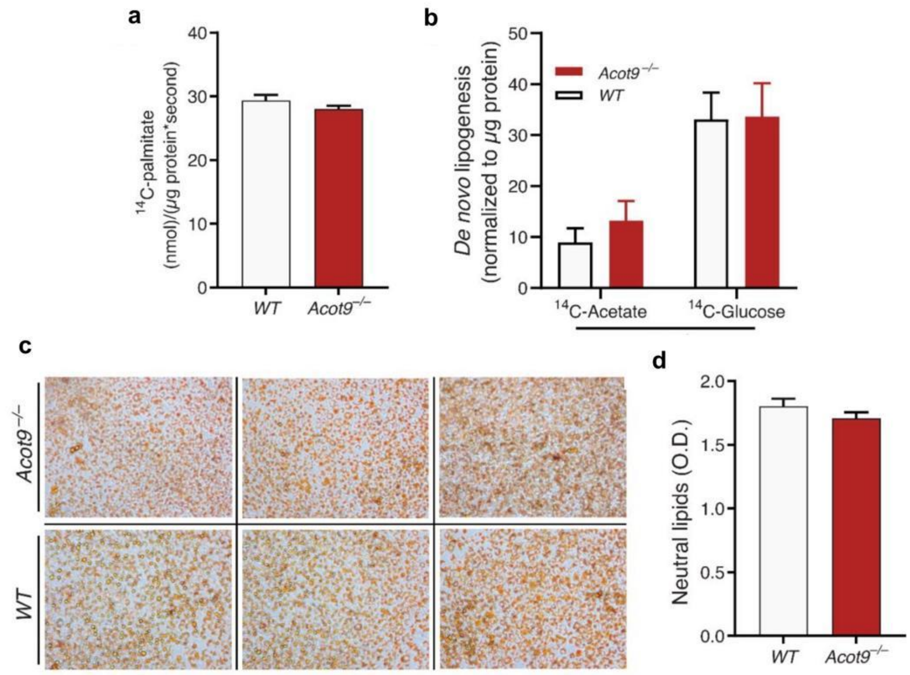


**Extended Data Fig. 14|Lipid uptake, synthesis, and storage in *Acot9*$^{-/-}$ and WT primary brown adipocytes.** **a**, Fatty acid uptake as measured by $^{14}$C-Palmitate uptake within 30 seconds of exposure. **b**, *De novo* lipogenesis rates were measured using $^{14}$C-acetate or $^{14}$C-glucose as substrate. **c**, Representative images of oil-red O-stained primary brown adipocytes from HFD-fed mice. **d**, Absorbance (492 nm) of oil-red O extracted from the primary brown adipocytes of HFD-fed mice. For a, b, and d, bars indicate the mean and error bars indicate + SEM; ns= not significantly different, Student's t-test.

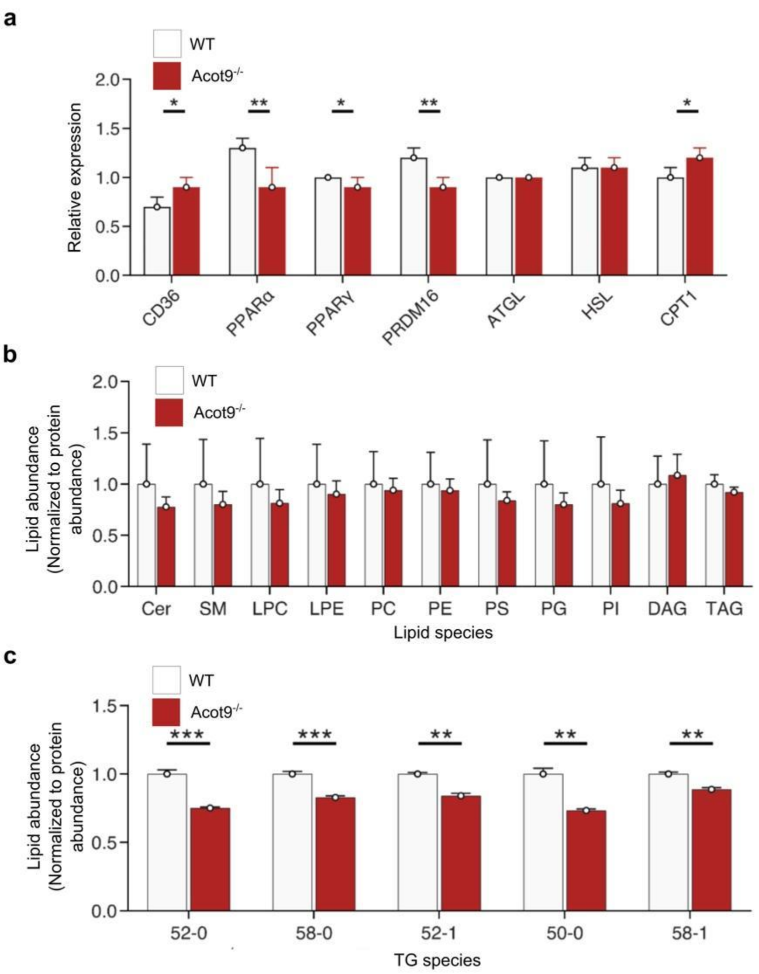


**Extended Data Fig. 15|Reduced levels of saturated and monounsaturated triacylglyceride (TG) species in *Acot9*$^{-/-}$ mice. a**, RT-qPCR analyses of metabolic genes in BAT of HFD-fed *Acot9*$^{-/-}$ and WT mice (n=5/genotype). Relative abundances were determined by normalizing to the mean of three housekeeping genes: Gapdh (glyceraldehyde 3-phosphate dehydrogenase), Tbp (TATA box binding protein), and Rpl32 (ribosomal protein L32). Bars indicate mean and error bars indicate + SEM. *P<0.05, **P<0.01, independent samples t-test. **b**, Mass spectrometry analyses of pooled lipid species and **c**, TG species that are statistically different between BATs of HFD-fed *Acot9*$^{-/-}$ and WT mice (n=4/genotype). For a and b, bars indicate the mean and error bars indicate + SEM; **P<0.01, ***P<0.00, independent samples t-test.

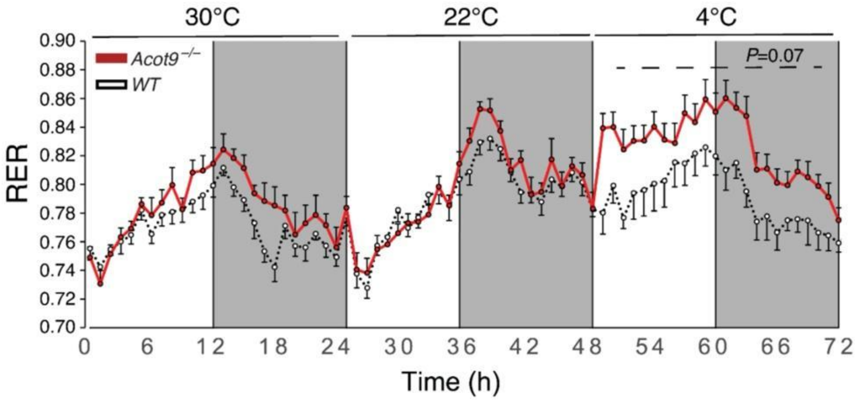


**Extended Data Fig. 16|Respiratory exchange ratio (RER) of HFD-fed *Acot9*$^{-/-}$ and WT mice.** Indirect calorimetric analysis of RER in HFD-fed *Acot9*$^{-/-}$ and WT mice (n=6/genotype). Graphs show mean ± SEM; analyzed by independent samples t-test at each time point.

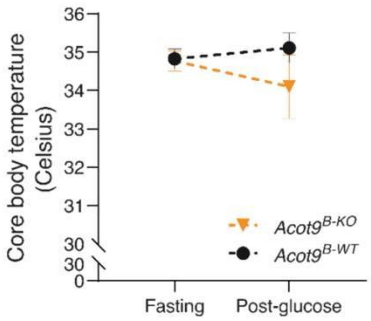


**Extended Data Fig. 17|Core body temperatures of *Acot9*$^{B\text{-}KO}$ and *Acot9*$^{B\text{-}WT}$ mice after glucose challenge.** Body temperatures of HFD-fed *Acot9*$^{B\text{-}KO}$ and *Acot9*$^{B\text{-}WT}$ mice (n=6/genotype) were analyzed by a rectal thermometer before and 30 m after a glucose challenge (2 g/kg body mass). Graphs show mean ± SEM; analyzed by independent samples t-test of the linear regression fit slope for each mouse.
